# BEATRICE: Bayesian Fine-mapping from Summary Data using Deep Variational Inference

**DOI:** 10.1101/2023.03.24.534116

**Authors:** Sayan Ghosal, Michael C. Schatz, Archana Venkataraman

## Abstract

We introduce a novel framework BEATRICE to identify putative causal variants from GWAS statistics. Identifying causal variants is challenging due to their sparsity and high correlation in the nearby regions. To account for these challenges, we rely on a hierarchical Bayesian model that imposes a binary concrete prior on the set of causal variants. We derive a variational algorithm for this fine-mapping problem by minimizing the KL divergence between an approximate density and the posterior probability distribution of the causal configurations. Correspondingly, we use a deep neural network as an inference machine to estimate the parameters of our proposal distribution. Our stochastic optimization procedure allows us to simultaneously sample from the space of causal configurations. We use these samples to compute the posterior inclusion probabilities and determine credible sets for each causal variant. We conduct a detailed simulation study to quantify the performance of our framework against two state-of-the-art baseline methods across different numbers of causal variants and different noise paradigms, as defined by the relative genetic contributions of causal and non-causal variants. We demonstrate that BEATRICE achieves uniformly better coverage with comparable power and set sizes, and that the performance gain increases with the number of causal variants. We also show the efficacy BEATRICE in finding causal variants from the GWAS study of Alzheimer’s disease. In comparison to the baselines, only BEATRICE can successfully find the APOE *ϵ*2 allele, a commonly associated variant of Alzheimer’s. Thus, we show that BEATRICE is a valuable tool to identify causal variants from eQTL and GWAS summary statistics across complex diseases and traits.

## 1. Introduction

Genome-Wide Association Studies (GWAS) provide a natural way to quantify the contribution each genetic variant to the observed phenotype [31]. However, the univariate nature of GWAS does not take into account the correlation structure shared between the genetic variants due to low recombination of nearby DNA regions [32]. Strong correlations can inflate the effect size of a non-causal genetic variant, thus leading to false positive identifications [3] Fine-mapping [23, 29] addresses this problem by analyzing the correlation structure of the data to identify small subsets of causal genetic variants [29, 27]. These subsets, known as credible sets, capture the uncertainty of finding the true causal variant within a highly correlated region [16]. Unlike p-values, the corresponding posterior inclusion probabilities (PIPs) computed during fine-mapping can be compared across studies of different sample sizes.

Traditional fine-mapping methods can be grouped into two general categories. The first category uses a penalized regression model to predict the output phenotype based on the collection of genetic variants [8, 26]. Popular regularizations like LASSO [30] and Elastic Net [26] simultaneously perform effect size estimation while slowly shrinking the smaller effect sizes to zero. The drawback of penalized regression models is that they optimize phenotypic prediction and, due to the correlation structure, do not always identify the true causal variants. The second category relies on Bayesian modeling. Here, the phenotype is modeled as a linear combination of the genetic variants, with sparsity incorporated into the prior distribution for the model weights. Approximate inference techniques, such as Markov Chain Monte Carlo (MCMC) [13] and variational methods [4] have been used to infer the effect sizes, PIPs, and credible sets. While these approaches represent valuable contributions to the field, they require the raw genotype and phenotype information, which raises privacy and regulatory concerns, particularly in the cases of publicly shared datasets. MCMC sampling also requires a burn-in period, which adds a substantial (100X) runtime overhead.

In response to these concerns, fine-mapping approaches have moved towards using summary statistics, which can be easily shared across sites. For example, the works of [2, 5, 15] use a stochastic or exhaustive search to identify the posterior probabilities of the causal configurations. However, exhaustive search based methods are restricted by the number of assumed causal variants, as this leads to an exponential increase in the dimensionality of the approximate posterior distribution. Stochastic search approaches [2] are less computationally expensive, but, by construction, they cannot handle infinitesimal effects from non-causal variants. Another fine-mapping approach is SuSiE [40], which estimates the variant effect sizes as a sum of “single effects”. These “single effect” vectors contain one non-zero element representing a causal variant and are estimated using a Bayesian step-wise selection approach. SuSiE provides a simple framework to robustly estimate PIPs and credible sets; however, there is limited evidence for its performance given the presence of infinitesimal genetic effects. Such scenarios can appear due to polygenicity of the trait, trans-interactions of variants, or varying correlation structure of the genomic region. Finally, the most recent fine-mapping method is CARMA [37]. Unlike previous methods, CARMA assumes a spike-slab prior over the effect sizes and uses a stochastic shotgun bases sampling approach for estimating posterior probabilities.

In this paper, we introduce BEATRICE, a novel framework for **B**ayesian fin**E**-mapping from summ**A**ry da**T**a using deep va**R**iational **I**nferen**CE**^1^. In contrast to prior work, we approximate the posterior distribution of the causal variants given the GWAS summary statistics as a binary concrete distribution [22, 17], whose parameters are estimated by a deep neural network. This unique formulation allows BEATRICE to use computationally efficient gradient-based optimization to minimize the KL divergence between the proposal binary concrete distribution and the posterior distribution of the causal variants. In addition, our unique optimization strategy samples a representative set of causal configurations in the process of minimizing the empirical KL divergence; these configurations can be used to obtain the PIPs and the credible sets. We compare our model with two state-of-the-art fine-mapping approaches, SuSiE [40] and FINEMAP [2]. We perform an extensive simulation study and quantify the performance of each model across increasing numbers of causal variants and increasing noise, as determined by the degree to which non-causal variants explain the phenotype variance. The runtimes of both SuSiE and BEATRICE are under one minute, which is significantly less than the runtimes of FINEMAP and CARMA. On average BEATRICE achieves a 2.2 fold increase in coverage, a 10% improvement in AUPRC, and similar power in comparison to SuSiE and FINEMAP.

## 2. Materials and Methods

### 2.1. Generative Assumptions of Fine-mapping

BEATRICE is based on a generative additive effect model. Formally, let **y** ∈ ℝ^*n*×1^ denote a vector of (scalar) quantitative traits across *n* subjects. The corresponding genotype data **X** ∈ ℝ^*n*×*m*^ is a matrix, where *m* represents the number of genetic variants in the analysis. Without loss of generality, we assume that the the columns of **X** have been normalized to have mean 0 and variance 1, i.e., 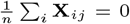 and 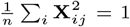 for *j* = 1, …, *m*. The quantitative trait is generated as follows:

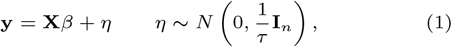

where *β* ∈ ℝ^*m*×1^ is the effect size, *η* ∈ ℝ^*n*×1^ is additive white Gaussian noise with variance 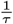, and **I**_*n*_ is the identity matrix.

### 2.2. Genome Wide Association Studies (GWAS)

GWAS uses a collection of element-wise linear regression models to estimate the effect of each genetic variant. Mathematically, the GWAS effect sizes are computed as 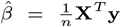, with the corresponding vector of normalized z-scores equal to 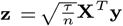 [31, 5]. The main drawback of GWAS is that non-causal genetic variants can have large effect sizes due to polygenicity of the quantitative trait [7], varying degrees of linkage disequilibrium (LD) with causal variants [3], and/or interactions of the variant with enriched genes [7]. One popular strategy to mitigate this drawback is to impose a sparse prior over *β* given the set of causal variants:

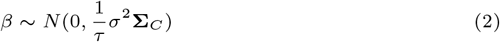

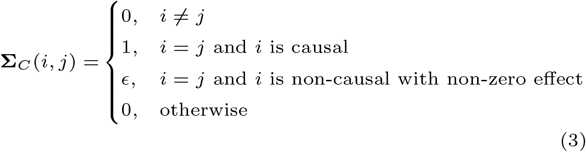

Notice from Eq. (3) that the variance of *β*(*i*) for a causal variant is 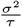 and the variance of *β*(*i*) for a non-causal variant with non-zero effect is 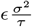, where *ϵ* is assumed to be small. This formulation handles residual influences from the non-causal variants, which are often observed in real-world data. Under this assumed prior, we can show [28, 5] that the normalized GWAS effect sizes **z** are distributed as:

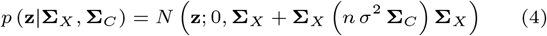

where 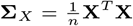 is the empirical correlation matrix of the genotype data, also known as the LD matrix. Broadly, the goal of fine-mapping is to identify the diagonal elements of Σ_*C*_ that corresponds to 1 given the effect sizes **z** and the LD matrix **Σ**_*X*_. The derivation is provided in **Section S1.1** of the Supplement.

### 2.3. The Deep Bayesian Variational Model

BEATRICE uses a variational inference framework for fine-mapping. For convenience, we represent the diagonal elements of **Σ**_*C*_ by the vector **c** ∈ ℝ^*m*×1^, where **c** encodes the causal variant locations. Given that we do not know the effect sizes and locations of the causal and non-causal variants, rather than fixing *ϵ* in Eq. (3), we assume that the diagonal elements of Σ_*C*_ are drawn from a binary concrete distribution, which can be viewed as a continuous relaxation of the Bernoulli distribution. Under this assumption, the variance of the effect size *β*_*i*_ is modeled as 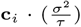, where **c**_*i*_ ∈ [0, 1] is a binary concrete random variable. The binary concrete distribution provides support for small non-zero values, which can be automatically learned from the data during inference, as described below.

Figure 1 provides an overview of BEATRICE. Our framework consists of three main components: an inference module, a random sampler, and a generative module. The inputs to BEATRICE are the summary statistics **z** and the LD matrix **Σ**_*X*_. The inference module estimates the parameters **p** of our proposal distribution *q* (*·*; **p**, *λ*) using a neural network. The random process sampler uses the parameters **p** to sample potential causal vectors **c** according to the given proposal distribution. Finally, the generative module calculates the likelihood of the observed summary statistics **z** via Eq. (4).

**Fig. 1.**
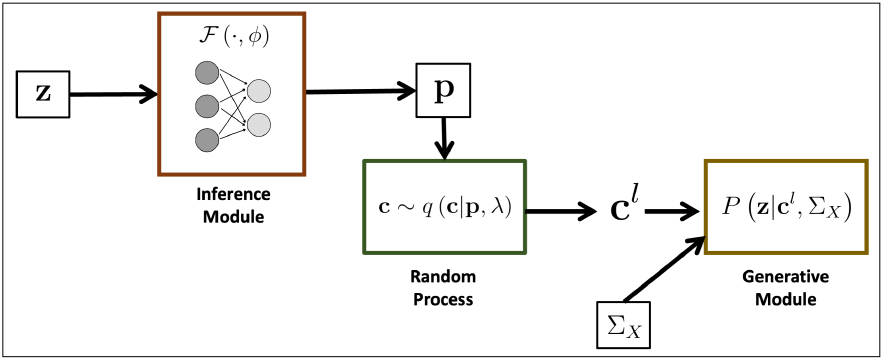
Overview of BEATRICE. The inputs to our framework are the LD matrix **Σ**_*X*_ and the summary statistics **z**. The inference module uses a neural network to estimate the underlying probability map **p**. The random process generates random samples **c**^*l*^ for the Monte Carlo integration in Eq. (12). Finally, the generative module calculates the likelihood of the summary statistics from the sample causal vectors **c**^*l*^.

#### 2.3.1. Proposal Distribution

The goal of fine-mapping is to infer the posterior distribution *p*(**c**|{**z, Σ**_**X**_}), where **c** corresponds to the diagonal elements of **Σ**_*C*_. Due to the prior formulation in Eqs. (2-3), solving for the true posterior distribution is computationally intractable, as it requires a combinatorial search over the possible causal configurations. Thus, we approximate the posterior distribution *p*(**c**|{**z, Σ**_**X**_}) with a binary concrete distribution *q*(**c**; **p**, *λ*) [22], where the parameters **p** of the distribution are functions of the inputs {**z, Σ**_**X**_}. Samples **c** generated under a binary concrete distribution can be viewed as continuous relaxations of independent Bernoulli random variables. This reparametrization [17] allows us to learn **p** from the data using standard gradient descent.

Formally, let **c**_*i*_ and **p**_*i*_ denote the *i*th element of the vectors **c** and **p**, respectively. Each **c**_*i*_ is independent and is distributed

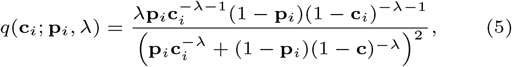

where the parameter *λ* controls the extent of relaxation from a Bernoulli distribution. We can easily sample from the binary concrete distribution in Eq. (5) via the relation

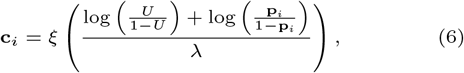

where *ξ*(*·*) is the sigmoid function, and the random variable *U* is sampled from a uniform distribution over the interval [0, 1]. Here, **p**_*i*_ specifies the underlying probability map and *U* provides stochasticity for the sampling procedure in Eq. (6). In practice, the gradient of Eq. (6) with respect to **p**_*i*_ tends to have a low variance, which helps to stabilize the optimization. The relationship between the random variable and the parameters are provided in Supplementary **Section S1.2**.

Intuitively, every element of the random vector **c** can be regarded as a continuous relaxation from a Bernoulli random variable, where *λ* controls the extent of relaxation from the 0/1 Bernoulli distribution. This continuous representation allows us to model the infinitesimal effects of the non-causal variants. Additionally, the underlying probability map **p** captures the relative importance of a variant containing a causal signal. The two unique properties of the probability maps are 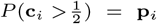 and lim_*λ*→0_ *P* (**c**_*i*_ = 1) = **p**_*i*_. The first property indicates that **p**_*i*_ controls the degree to which **c**_*i*_ assumes low values close to 0 and high values close to 1. This property also give BEATRICE flexibility to handle genetic variants with different levels of association, thus aligning with our generative process that assumes some non-causal variants may have small non-zero effects. The second property implies that a high probability **p**_*i*_ at location *i* is highly indicative of a causal variant. Taken together, the binary concrete distribution has an easily-optimized parameterization with desirable properties.

#### 2.3.2. Variational Inference

We select the variational parameters {**p**, *λ*} to minimize the Kullback–Leibler (KL) divergence between the proposal distribution and the posterior distribution of the causal vector **c** given the input data {**z, Σ**_**X**_}, that is

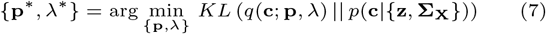

Using Bayes’ Rule, Eq. (7) can be rewritten as follows:

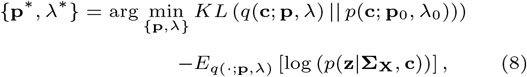

where we have assumed an element-wise binary concrete prior *p*(**c**; **p**_0_, *λ*_0_) over the vector **c**. We fix the relaxation parameter to be small (*λ* = 0.01) and the probability map to be uniform 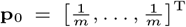. Thus, the first term of Eq. (8) can be viewed as a regularizer that encourages sparsity in causal vectors **c**. The second term of Eq. (8) can be interpreted as the likelihood of the observed test statistics. The works of [25, 33] have demonstrated that under certain assumptions, the likelihood term of the summary statistics is the same as the original data likelihood *p* (**y**|**X, c**) derived from Eq. (1).

During optimization, the relaxation parameter *λ* is annealed [22, 17] to a small non-zero value (0.01) with fixed constant rate, and the underlying probability map **p** is optimized using gradient descent. Specifically, we use a neural network to generate the vector **p** = ℱ (**z**; *ϕ*). The details of the neural network architecture are provided in **Section S1.4** of the Supplement. Practically speaking, the neural network ties the input data {**z**, Σ_*X*_} to the parameter space of the proposal distribution in a data-driven fashion. Empirically, we find that generating **p** as a function of the input data regularizes the model and leads to a stable optimization.

Optimizing **p**\* now amounts to learning the parameters of the neural network *ϕ*. Given a fixed value of *λ*, the neural network loss function follows from Eq. (8) as

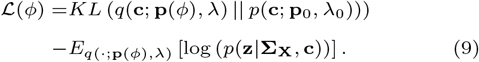

We have defined **p**(*ϕ*) ≜ ℱ (**z**; *ϕ*) for notational convenience.

#### 2.3.3. Optimization Strategy

Since Eq. (9) does not have closed-form expressions, we use Monte Carlo integration to accurately approximate *L*(*ϕ*) in the regime of small *λ*, i.e., when the binary concrete distribution behaves similarly to a Bernoulli distribution.

Let **c**^1^(*ϕ*), …, **c**^*L*^(*ϕ*) be a collection of causal vectors sampled independently from *q*(*·*|**p**(*ϕ*), *λ*) according to Eq. (6). The likelihood term of Eq. (9) is computed as

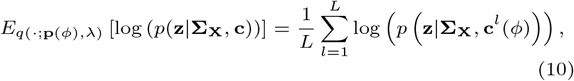

where the right-hand side probability is computed using Eq. (4) by substituting **c**^*l*^(*ϕ*) for the diagonal entries of **Σ**_*C*_ in each term of the summation. Once again, the continuous relaxation used to generate **c**^*l*^(*ϕ*) in Eq. (6) allows us to directly optimize *ϕ*.

We approximate the first term of Eq. (9) under the assumption of small {*λ, λ*_0_}. In this case, the binary concrete distribution behaves like a {0, 1} Bernoulli distribution. Under these conditions, we can write the first term of Eq. (9) as

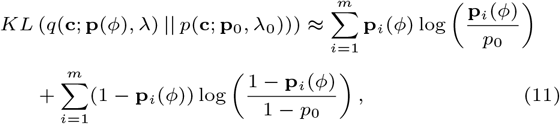

where *p*_0_ is a fixed scalar parameter used to construct the prior vector **p**_0_. We note that the criteria {*λ* → 0.01, *λ*_0_ = 0.01} is satisfied in practice, as *λ* is annealed during the optimization to progressively smaller values and *λ*_0_ is fixed *a priori*. This approximations allow us to rewrite the neural network loss as

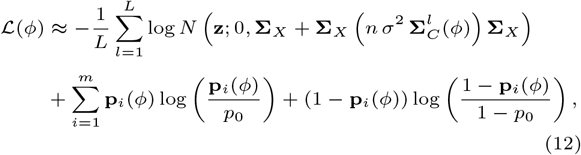

where 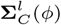 is a diagonal matrix with **c**^*l*^(*ϕ*) as the diagonal entries. We use a stochastic gradient descent optimizer [18] to minimize the loss ℒ (*ϕ*) with respect to the neural network weights *ϕ*. This process is detailed in Algorithm 1.

In our optimization process, we further take advantage of the sparsity of the causal vector, leading to significant improvements in computational complexity. The complexity analysis is provided in the **Section S1.5**.

## 3. Verification and Comparison

### 3.1. Causal Configurations and PIPs

We evaluate Posterior Inclusion Probabilities (PIPs) and credible sets of each method. PIPs estimate how likely each variant is causal as a measure of its importance. Credible sets are subsets of variants that likely contain a causal variant, which captures the uncertainty of finding the true variant.

#### Algorithm 1 Optimization scheme to minimize Eq. (12)

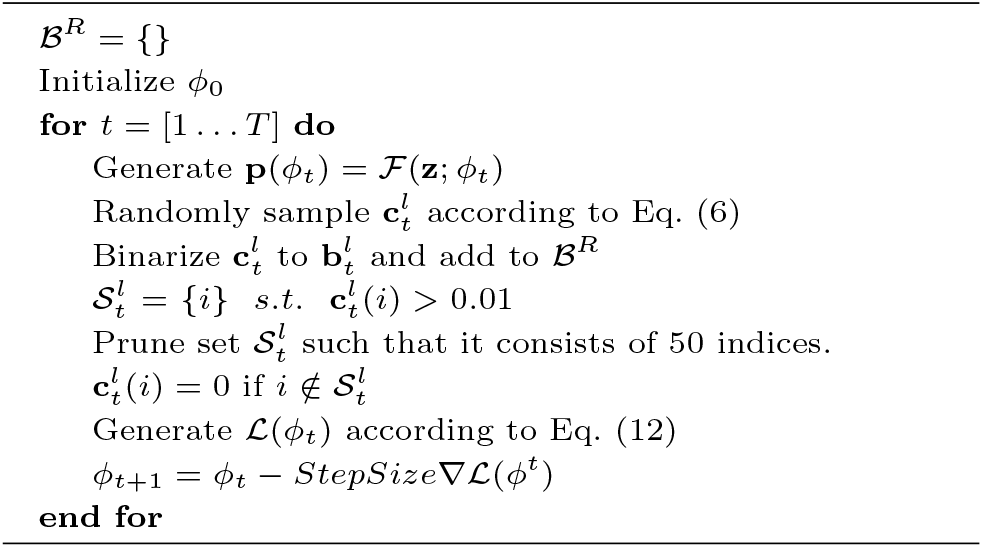

The main challenge to estimating the posterior probability of a given causal configuration (i.e., set of causal variant locations) is the exponentially large search space. Let **b** denote a binary vector with a value of 1 at causal locations and a value of 0 at non-causal locations. At a high level, **b** can be viewed as a binarized version of the causal vector **c** in the previous sections. Using Bayes’ Rule, the posterior probability of **b** given the input data {**z, Σ**_**X**_} can be written as follows:

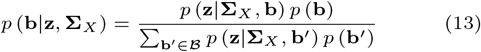

where ℬ is the set of all 2^*m*^ possible causal configurations, **z** denotes the summary statistics, and **Σ**_*X*_ is the LD matrix. Even though ℬ is exponentially large, it has been argued [14] that the majority of these configurations have negligible probability and do not contribute to the denominator of Eq. (13).

Our stochastic optimization provides a natural means to track causal configurations with non-negligible probability in *p* (**b**|**z, Σ**_*X*_). Namely, at each iteration of stochastic gradient descent, we randomly generate a sample causal vector **c**^*l*^ to minimize Eq. (12). In parallel, we binarize the vector **c**^*l*^ via

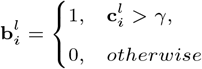

and add the resulting vector **b**^*l*^ to a reduced set of causal configurations ℬ^*R*^. The variational objective ensures that our proposal distribution converges to the true posterior distribution of the causal vectors. Thus, the samples **c**^*l*^ lie near modes of the posterior distribution, which can also be viewed as the neighborhood of non-negligible probability.

Our experiments use a threshold *γ* = 0.1 to binarize the vectors **c**^*l*^. The threshold *γ* can be viewed as a user-specified lower bound for *ϵ*, where fixing *γ* = 0.1 preserves only variants with an estimated effect size variance greater than 0.1*σ*^2^. Interestingly, the threshold *γ* can also be viewed as a sparsity penalty that loosely controls the size of the reduced set of causal configurations ℬ^*R*^. For example, when *γ* is large, we include only a small set of causal configurations with high posterior probability, whereas when *γ* is small, we allow for more configurations to be included in our analysis. A higher threshold is often beneficial in the presence of large interaction effects from non-causal variants, and a lower threshold is useful when the causal variants are weakly associated with the outcome. Finally, we note that *γ* is also linked to the computational overhead of BEATRICE. When *γ* is large we will need fewer samples to estimate posterior configuration, as compared to smaller values of when *γ*, leading to lower time complexity. While we fix the default value for *γ* at 0.1, this parameter can be adjusted by the user as desired. In **Section S1.10** the sensitivity of the results with different values of *γ*.

After obtaining the sampled vectors, we replace the exhaustive set **B** in Eq. (13) with the reduced set ℬ^*R*^ for tractable computation of *p* (**b**|**z, Σ**_*X*_). We then compute the posterior inclusion probability (PIP) of each variant by summing the probabilities over the subset of ℬ^*R*^ with a value of 1 at that variant location. Mathematically,

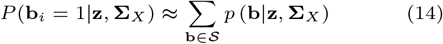

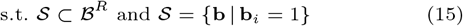

where 𝒮 is a subset of ℬ^*R*^ that contains a 1 at location *i*.

Finally, we identify the credible sets in two steps. First, in a conditional step-wise fashion, we identify the variants with the highest conditional probability given the previously selected variants. This strategy identifies the set of “key” variants with a high probability of being causal. Second, we determine the credible set for each key variant, by computing the conditional inclusion probabilities of each variant given the key variants and adding variants to the credible set. A detailed description of this process can be found in the Supplementary Methods document (**Section S1.3 in S1 text**).

### 3.2. Baselines

We compare our approach with these state-of-the-art methods:

- **Finemap:** This approach [2] uses a stochastic shotgun search to estimate the PIPs and the credible sets.
- **SuSiE:** [40, 34] introduced an iterative Bayesian selection approach for fine-mapping that represents the causal vector as a sum of “single-effect” vectors.
- **CARMA:** The work of [37] introduced a Bayesian approach for fine-mapping using a spike-and-slab prior over the effect sizes to model the GWAS summary statistics.

Further details are provided in Supplementary Section S1.15.

### 3.3. Evaluation Strategy

We evaluate several metrics of performance for each method.

#### Evaluating PIPs

We compare the quality of the PIPs via the AUPRC metric. AUPRC (area under the precision-recall curve) is computed by sweeping a threshold on the PIPs and computing precision and recall against the true configuration of causal and non-causal variants. High precision indicates a low false positive rate in the estimated causal variants. High recall indicates that the model correctly identifies more of the causal variants. Thus, the AUPRC, can be viewed as a holistic measure of performance across both classes. AUPRC is also robust to severe class imbalance [10], which is the case in fine-mapping, as the number of causal variants is small. Additionally, in Supplementary Section S1.8 we visualize power vs. FDR for different thresholds of the PIPs. Following standard nomenclature, both power and recall measures the probability of detection 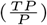 and FDR (1-precision) measures the type-I error 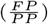. Therefore, the power versus FDR curve provides a visual comparison, while AUPRC gives the numeric quantification of the performances.

#### Coverage, Power and Size of the Credible Sets

We follow the strategy of [40, 34] to define a credible set. A credible set is defined as a collection of variants that contain a single causal variant with a probability equal to the coverage. Given that the number of causal variants can be arbitrary, we use two metrics to assess the quality of the credible sets: coverage and power. Coverage is the proportion of credible sets that contain a causal variant, and power is the proportion of causal variants identified by *all* the credible sets. Higher coverage indicates that the method is confident about its prediction of *each* causal variant, whereas higher power indicates the method can accurately identify all the causal variants.

One caveat is that a method can generally achieve both higher coverage and higher power simply by adding variants to the credible sets. To counter this trend, we report the average size of the credible sets identified by each method. Ideally, we would like the credible sets to be as small as possible while retaining high coverage and high power.

## 4. Experimental Results

### 4.1. Setup for Simulation Experiments

#### Genotype Simulations

We use the method of [11] to simulate genotypes **X** based on data from the 1000 Genomes Project.

We select an arbitrary sub-region (39.9*Mb* − 40.9*Mb*) from Chromosome 2 as the base. After removing rare variants (MAF < 0.02), the remaining 3.5K variants are used to simulate pairs of haplotypes to generate 10,000 unrelated individuals. We do not run any filtering of variants on our simulated data. We chose a MAF threshold of 0.02, as it lies in the middle of the range 0.01 − 0.05 used in GWAS studies [1]. In each experiment below, we randomly select *m* = 1000 variants and *n* = 5000 individuals to generate the phenotype data.

#### Phenotype Generation

We generate the phenotype **y** from a standard mixed linear model [25], where the influences of the causal variants are modeled as fixed effects, and the influences of other non-causal variants are modeled as random effects. In this case, the genetic risk for a trait is spread over the entire dataset, with each variant having small individual effects, as per the polygenicity assumption of a complex trait. We randomly select the causal variants in our simulations. Thus, some simulations will have causal variants in LD, while others will select causal variants with low correlation.

Given a set of *d* causal variants *C*, let **X**_*C*_ ∈ **R**^*n*×*d*^ denote the corresponding subset of the genotype data and **X**_*NC*_ ∈ **R**^*n*×*m*−*d*^ denote the remaining non-causal variants. From here, we generate the phenotype data **y** as follows:

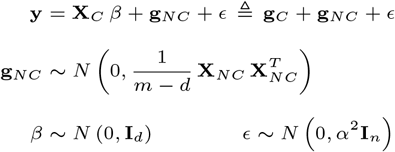

where *β* is the *d*-dimensional effect sizes sampled from a Gaussian distribution, and *ϵ* is an zero-mean Gaussian noise with variance *α*^2^. The random variable **g**_*NC*_ models the effect of the non-causal variants as a multivariate Gaussian vector with mean 0 and covariance 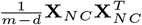. Likewise, **g**_*C*_ = **X**_*C*_ *β* captures the effect of the causal variants.

In our experiments, we define *ω*^2^ as the total phenotypic variance attributed to the genotype (both **g**_*C*_ and **g**_*NC*_) and *p* as the proportion of this variance associated with the causal variants in **g**_*C*_. Using the strategy described in [24], we enforce these conditions by normalizing the phenotype **y** as follows:

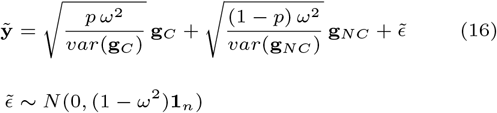

where *var*(**g**_*C*_) and *var*(**g**_*NC*_) are the empirical variances of the genotype vectors **g**_*C*_ and **g**_*NC*_, respectively.

To replicate real-world scenarios, we use a GWAS setup to estimate the effect size 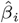 of each variant *i* based on the phenotype 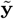 and the un-normalized genotype data 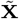. Here, we use a simple linear model and ordinary least squares to estimate the effect sizes. From here, we convert the estimated effect sizes to z-scores via 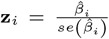, where *se*(*·*) denotes the standard error. The LD matrix is computed as 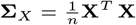, where **X** is the normalized genotype data. The z-scores and LD matrix are input to each of the fine-mapping methods above.

#### Noise Configurations

We evaluate the performance of each method while varying the number of causal variants *d*, the total genotype variance *ω*^2^, and the proportion of this variance associated with the causal variants *p*. Formally, we sweep over the ranges *d* = [1, 4, 8, 12], *ω*^2^ = [0.1, 0.2, 0.4, 0.5, 0.7, 0.8], and *p* = [0.1, 0.3, 0.5, 0.7, 0.9]. For each parameter setting, we randomly generate 20 datasets by independently re-sampling the causal variant locations, the effect sizes {*β*_*i*_}, the non-causal component **g**_*NC*_, and the noise 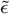. We run all fine-mapping methods over a total of 4 × 6 × 5 × 20 = 2400 configurations.

### 4.2. Application to Real-World SNP Data

We compare the performance of each finemapping method on a GWAS study of Alzheimer’s Disease (AD). AD is a polygenic disorder, making it an ideal test bed to evaluate each model. We use the publicly available GWAS summary statistics released by [36] to obtain the z-scores and the UK Biobank data to generate the LD matrices. We compare the finemapping results of SuSiE and BEATRICE. Notice that we cannot run FINEMAP because the reported GWAS statistics only contain the z-scores, whereas FINEMAP requires the effect sizes 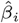 and the corresponding standard errors to perform finemapping. This real-world study also highlights a major drawback of FINEMAP, should the original effect sizes and standard errors be unavailable. Both SuSiE and BEATRICE were run with their default setting.

#### Data Acquisition

The recent GWAS analysis performed by [36] identified multiple statistically significant index SNPs in AD. We clumped the GWAS statistics to 175 regions and finemapped the top 20 regions sorted by the p-values of the index SNPs. Additionally, we sub-select SNPs from the GWAS summary statistics that overlap with both the publicly available 1000 Genome Phase-3 data and the UK Biobank data.

#### Data Preprocessing

We first filter out variants that are not common to both the 1000 Genome database and the UK Biobank database. We further removed the strand ambiguous SNPs (i.e., those with complementary alleles, either C/G or A/T SNPs) due to the lack of strand information. For example, an A/T allele can be mapped to T/A, or A/T based on the strand information. Without this information, we cannot map the allele in the base data and GWAS results. After filtering, we verify that the counted alleles are present in the GWAS summary statistics for AD and the UK biobank LD matrices. We identify 175 clumps using the GWAS statistics, where each clump contains SNPs less than 250KB away from the index SNP with *R*^2^ > 0.1 and *MAF* > 0.01. We note that the GWAS statistics published by [36] reported multiple SNPs in the first clump with z-scores of infinity. For numerical stability, we clip these z-scores to 200. Finally, we merge overlapping clumped regions into a single region. Further details about the clumped variants are reported in Supplementary Table 1. The resulting variants within each clump are used to generate LD matrices using the publicly available ^2^ LD matrices derived from 337K subjects of the UK Biobank database [35].

### 4.3. Model Performance

#### 4.3.1. Simulated Experiments

##### Varying the Number of Causal Variants

Figure 2 illustrates the performance of each method (BEATRICE, FINEMAP, SuSiE, and CARMA) while increasing the number of causal variants from *d* = 1 to *d* = 12. The points denote the mean performance across all noise configurations (*ω*^2^, *p*) for fixed *d*, and the error bars represent the 95% confidence interval across these configurations. We note that BEATRICE achieves a uniformly higher AUPRC than all baseline methods, which suggests that BEATRICE can better estimate the PIPs than CARMA, FINEMAP or SuSiE. BEATRICE also provides a 0.9 − 1.4-fold increase in coverage than the baselines while maintaining a similar power, which indicates that the credible sets generated by BEATRICE are more likely to contain a causal variant than the baselines. Finally, we note that BEATRICE, identifies the same or smaller credible set sizes than CARMA, FINEMAP, and SuSiE. Taken together, as the number of causal variants increases, BEATRICE gives us a better estimate of the PIPs. Unlike the baseline methods, BEATRICE does not impose any prior assumptions over the total number of causal variants, which may lead to its improved performance. Finally, we observe that CARMA achieves notably lower AUPRC and power than all other methods and is comparable to SuSiE and FINEMAP in the other metrics.

**Fig. 2.**
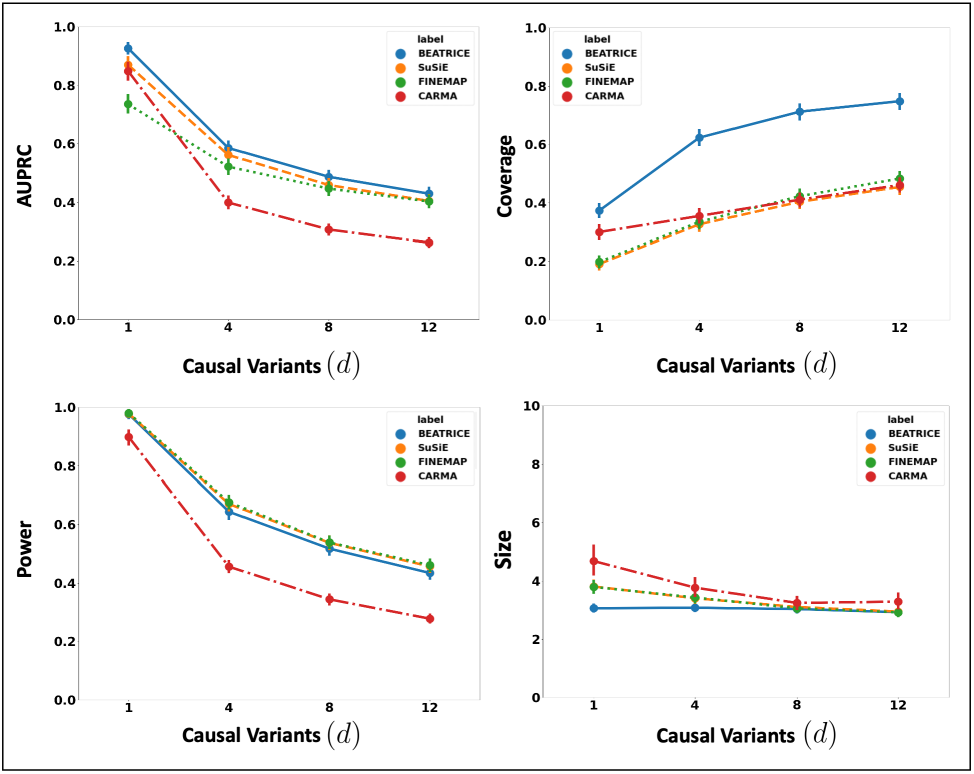
The performance metrics for the three methods across varying numbers of causal variants. Along the x-axis, we plot the number of causal variants, and across the y-axis, we plot the mean and confidence interval (95%) of each metric. We calculate the mean by fixing *d* to a specific value *d* = *d*\* and sweep over all the noise settings where *d* = *d*\*.

##### Increasing the Genotype Contribution

Figure 3 shows the performance of each method while increasing the genetically-explained variance from *ω*^2^ = 0.1 to *ω*^2^ = 0.8. Similar to the above experiment, the points in the figure denote the mean performance across all other noise configurations (*d, p*) for fixed *ω*^2^, and the error bars represent the 95% confidence intervals across these configurations. We note that BEATRICE achieves a significantly higher AUPRC than FINEMAP and CARMA and a slightly higher AUPRC than SuSiE. When evaluating the credible sets, we observe similar trends in coverage (BEATRICE is 0.25 − 2.34 folds higher) and power (similar performance across methods). Once again, CARMA achieves significantly lower AUPRC and power. All four methods identify credible sets of similar size. We submit that BEATRICE achieves the best trade-off across the four performance metrics.

**Fig. 3.**
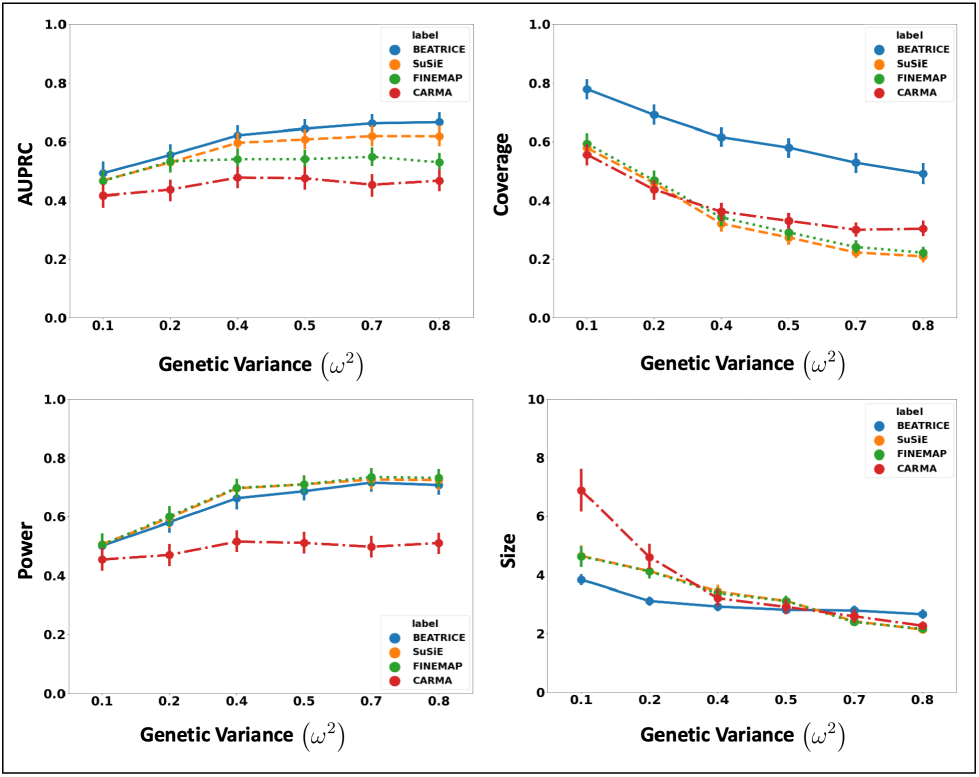
The performance metric for increasing phenotype variance explained by genetics. Along the x-axis, we plot the variance explained by genetics (*ω*^2^), and across the y-axis, we plot each metric’s mean and confidence interval (95%). We calculate the mean by fixing *ω*^2^ to a specific value *ω* = *ω*\* and sweep over all the noise settings where *ω* = *ω*\*.

##### Varying the Contributions of Causal and Non-Causal Variants

Figure 4 illustrates the performance of each method while increasing the contribution of the causal variants from *p* = 0.1 to *p* = 0.9. Again, the points denote the mean performance across all other noise configurations (*d, ω*^2^) for fixed *p*, and the error bars represent the 95% confidence intervals across these configurations. From an application standpoint, the presence of non-causal variants with small non-zero effects makes it difficult to detect the true causal variants. Accordingly, we observe a performance boost across all methods when *p* is larger. Similar to our previous experiments, BEATRICE provides the best AUPRC, with converging performance as *p* → 1. In addition, BEATRICE identifies smaller credible sets with significantly higher coverage while maintaining power. Thus, we conclude that BEATRICE is the most robust of the three methods to the presence of noise from non-causal variants. This performance gain may arise from our binary concrete proposal distribution for the causal vector **c**, which provides flexibility to accommodate varying degrees of association. Finally, we note that compared to all the approaches, CARMA has significantly lower AUPRC and power, which suggest that it fails to identify the true causal SNPs across the various noise paradigms. Moreover, as shown in Figure 10, the runtime of CARMA is significantly higher than the others due to its MCMC procedure. Thus, we omit it from our subsequent analyses.

**Fig. 4.**
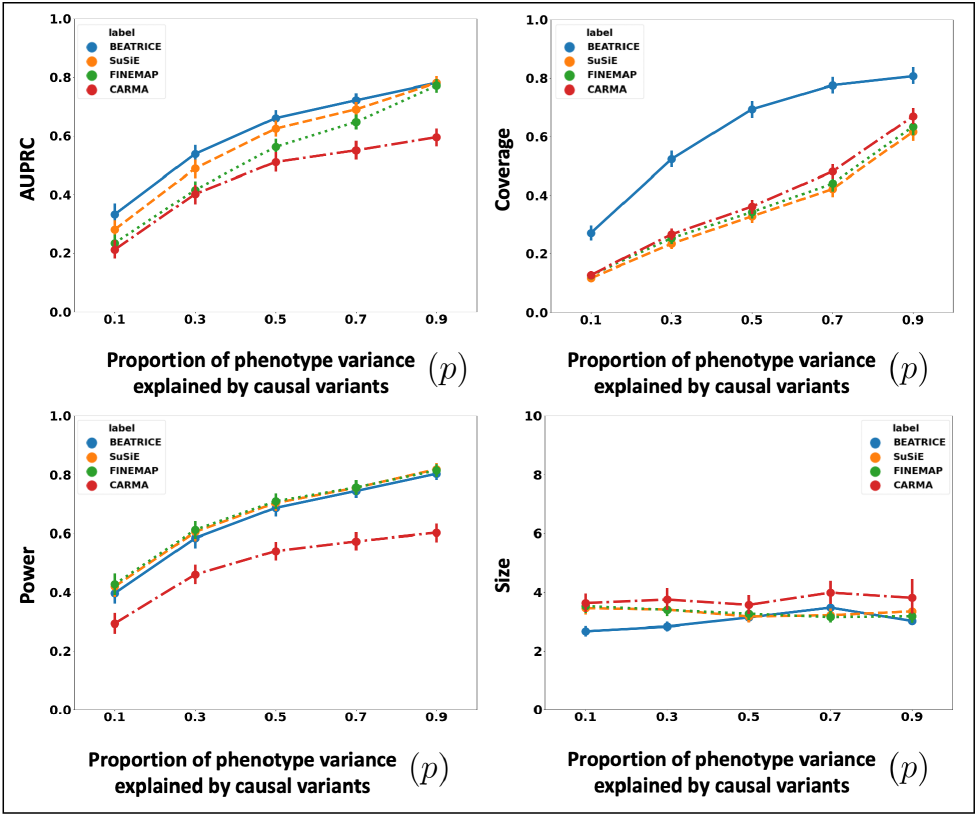
The performance metric for multiple levels of noise introduced by non-causal variants. The noise level (*p*) is explained by the variance ratio of non-causal variants vs. causal variants. Along the x-axis, we plot the noise level (*p*); across the y-axis, we plot each metric’s mean and confidence interval (95%). We calculate the mean by fixing *p* to a specific value *p* = *p*\* and sweep over all the noise settings where *p* = *p*\*.

#### 4.3.2. Finemapping Results on Real-World AD Data

Figure 5 shows the absolute z-scores of the SNPs in each of the 20 clumps. The GWAS statistics of [36] report multiple z-scores with high values in each clump. The inflation in the GWAS statistics could result from the infinitesimal effects of multiple SNPs within each clump. This scenario is similar to the simulation settings when *p* is small.

**Fig. 5.**
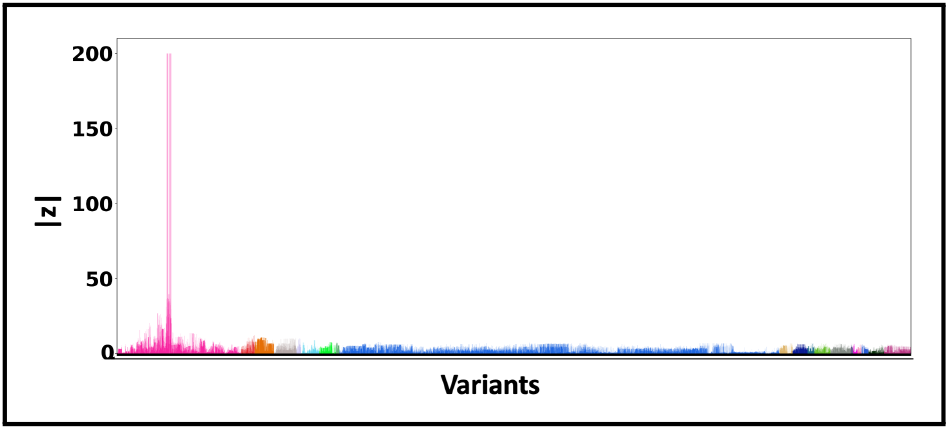
Z-scores of the variants present each of the 3 clumps. The scores are obtained from a GWAS study of Alzheimer’s Disease [36] colored by locus. The x-axis denotes the SNP index present in each locus. The y-axis reports the absolute z-scores.

Figure 6 shows the PIPs (> 0.9) identified by each finemapping method, colored by clump (i.e., locus). The x-axis corresponds to the SNP index, and the y-axis reports the corresponding PIP values. We explore the variants from the first clump (chr19:44515074-46457976), which contains the TOMM40, APOC1, and APOE genes. These genes have been repeatedly identified as potential disease-causing loci for AD [21, 6, 38, 9]. In Supplementary Tables 2 and 3 we report the SNPs with PIP> 0.9 for BEATRICE and SuSiE, respectively. Fig. 7 provides a detailed view of the z-scores, PIPs estimated by BEATRICE and SuSiE, and the LD structure of the variants within this clump. Interestingly, one of the high PIP SNP identified by BEATRICE is *rs7412*, the Apolipoprotein E (APOE) *ϵ*2 allele, which is a common landmark “risk” factor for Alzheimer’s disease [19]. As a second point of reference, Fig. 8 provides a detailed view of the variants in the ninth clump. Clump 9 (chr6:32123639-32900787) is the largest clump (≈ 8000 SNPs) in our analyses and overlies the HLA region, which is commonly known for complex LD [12] structure.

**Fig. 6.**
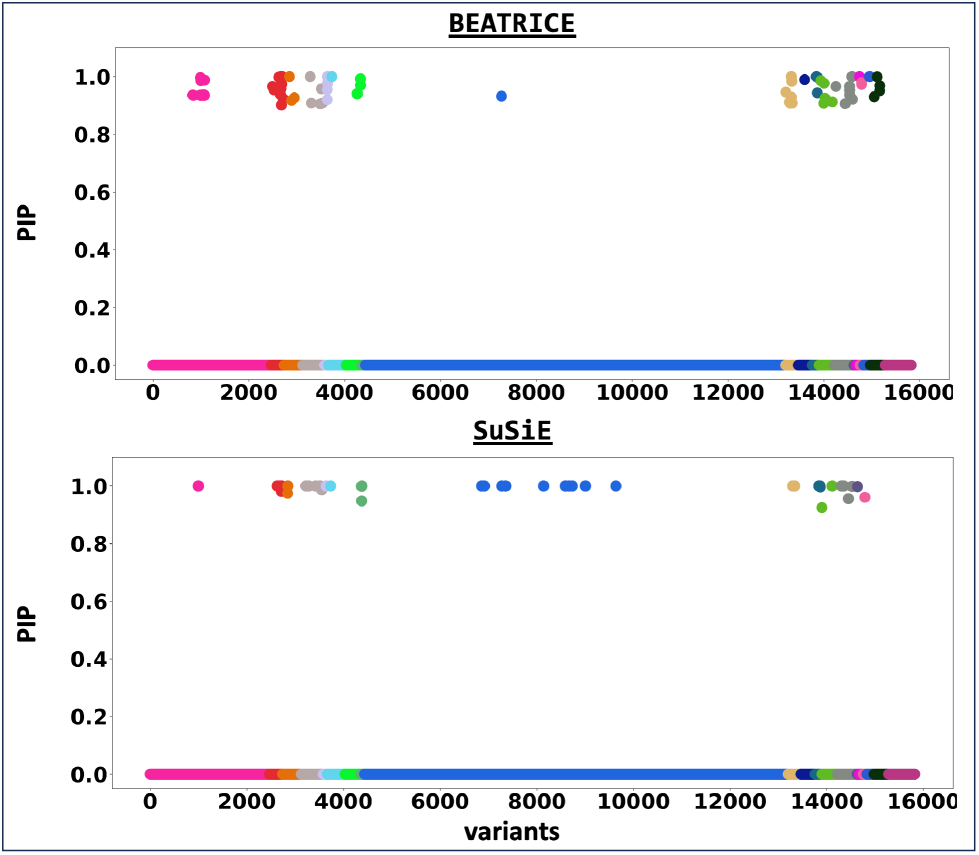
The posterior inclusion probabilities obtained by running finemapping on the top-20 clumps from the Alzheimer’s Disease GWAS. We report the PIPs of SNPs with PIP > 0.9. The x-axis denotes the SNP index present in each locus. The y-axis reports the PIP values.

**Fig. 7.**
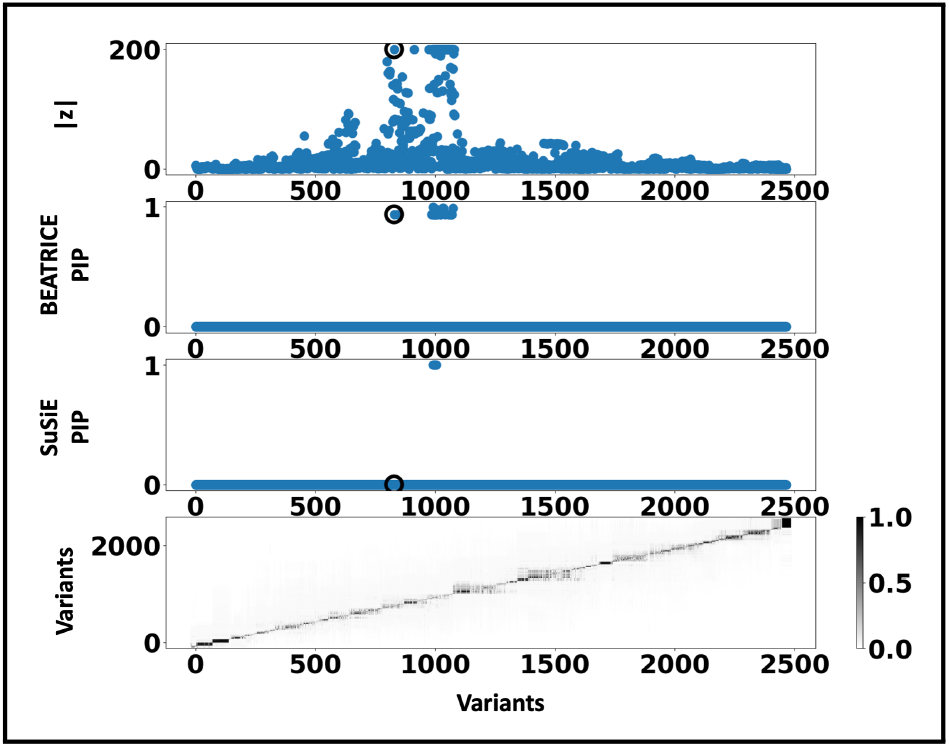
Detailed results for Clump 1: **Top Row:** Absolute z-scores of the included variants; the index SNP is indicated by a black circle. **Second Row:** PIPs identified by BEATRICE. **Third Row:** PIPs identified by SuSiE. **Bottom Row:** LD structure between the variants, where the colorbar indicates the *r*^2^ values.

**Fig. 8.**
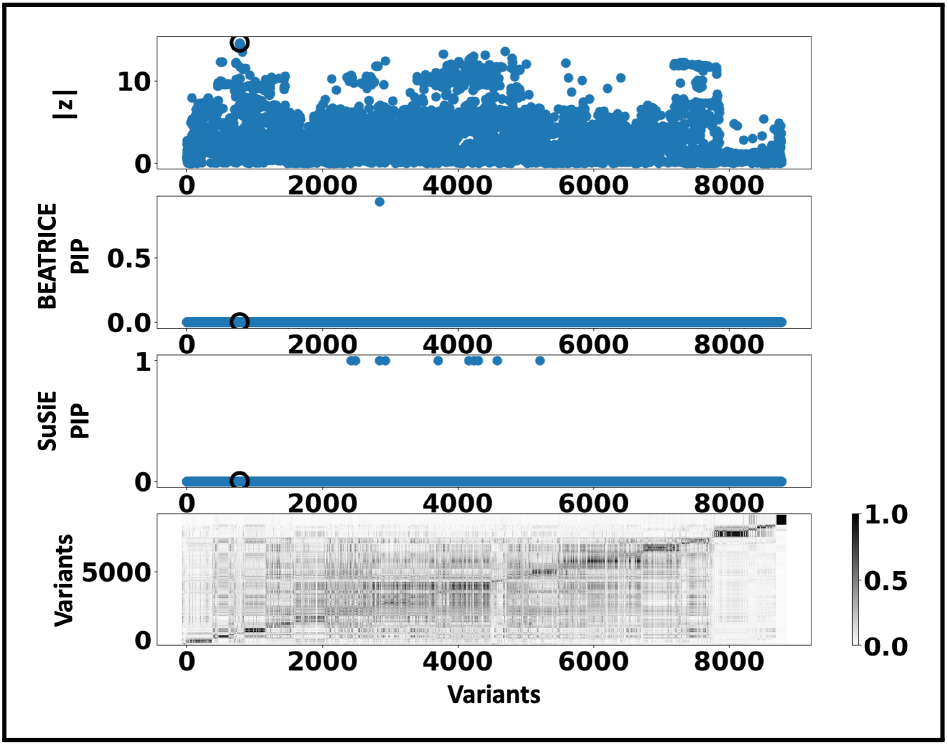
Detailed results for Clump 9: **Top Row:** Absolute z-scores of the included variants; the index SNP is indicated by a black circle. **Second Row:** PIPs identified by BEATRICE. **Third Row:** PIPs identified by SuSiE. **Bottom Row:** LD structure between the variants, where the colorbar indicates the *r*^2^ values.

In an exploratory analysis, we investigate the biological consequences of the SNPs with high PIPs (> 0.9) of the first clump, as identified by each method. Further details about this analysis are presented in Supplementary **Section S1.17**.

## 5. Discussion and Summary

BEATRICE is a novel, and general purpose tool for fine-mapping that can be used across a variety of studies. One key contribution of BEATRICE over methods like FINEMAP and SuSiE is its ability to discern infinitesimal effects from non-causal variants, including those in high LD with true causal variants. Our simulated experiments in **Section 4.3.1** and **Section S1.8** capture this improved performance by sweeping the proportion of the observed variance attributed to causal (fixed effects) and non-causal (random effects) genetic variants.

This parameter *p* ∈ [0, 1] is swept over its natural domain, such that *p* = 1 implies that the only link between the genotype and phenotype comes from the causal variants. At this extreme, all methods achieve comparable performance (Figure 4). However, as *p* decreases, meaning that the effects of non-causal variants increase, BEATRICE outperforms both baselines. In Figure 9 we show an example finemapping using BEATRICE for a simulation setting {*d* = 1, *ω*^2^ = 0.1, *p* = 0.3}. Notice that the z-score for the causal SNP is not the largest, which accurately highlights the importance of fine mapping. In Supplementary Section S6, we compare examples on the effects of SNP heritability explained by the non-causal SNPs. We show that when non-causal SNPs explain most of the genetic heritability, only BEATRICE can successfully assign the highest PIP to the causal SNPs. This result shows that BEATRICE can successfully use the binary concrete distribution to model non-causal variants with non-zero effects while the sparsity term of ℒ (*·*) prioritizes potentially causal variants.

**Fig. 9.**
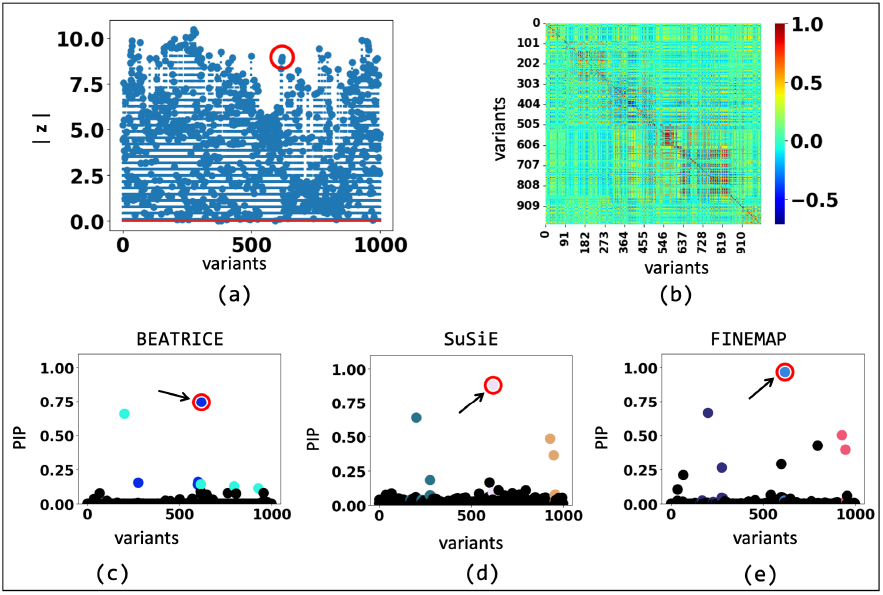
The fine-mapping performance of BEATRICE, SuSiE, and FINEMAP at a noise setting of {*d* = 1, *ω*^2^ = 0.1, *p* = 0.3}. (a) The absolute z-score of each variant as obtained from GWAS. (b) Pairwise correlation between the variants. (c)-(e) illustrate the posterior inclusion probabilities of each variant, as estimated by the three method. The red circle marked by an arrow shows the location of the causal variant. The non-black markers represent the variants assigned to a credible set, color-coded based on the assignment.

A second contribution of BEATRICE is our strategic integration of neural networks within a larger statistical framework. Specifically, we use the neural network in Figure S2 as an inference engine to estimate the parameters **p** of our proposal distribution. Effectively, we leverage the neural network as a universal function approximator to establish the relationship between the parameter space and the input data space. We choose neural networks as our inference engine over other types of functions due to their flexibility, scalability, and ease of optimization via backpropagation. We demonstrate in **Sections 4.3.1, S1.8, and S1.9** that the deep neural network can successfully generate sample causal configurations that well approximate the true posterior distribution leading to improved AUPR, power, coverage, and FDR. Moreover, BEATRICE leverages the continuous representation of the causal vectors **c**^*l*^ to backpropagate the gradients through the random sampler and train the network. These continuous representations of **c**^*l*^ result in low-variance gradients with respect to the underlying probability map, thus leading to a stable optimization.

Related to the above point, a third contribution of BEATRICE is its ability to identify the representative sets of causal configurations from the exponential search space to compute the PIPs and credible sets. In **Section S1.11**,**S1.12**, and **S1.14**, we show that BEATRICE generates well-calibrated PIPs in the presence of model misspecification. The improved performance can be attributed to our random sampling process, which ensures that the randomly sampled causal vectors slowly converge to the causal configurations with non-negligible posterior probability. Furthermore, this strategy allows us to efficiently estimate the PIPs in finite run-time. However, we note that the current implementation of BEATRICE does not estimate the posterior variant effect sizes. Instead, BEATRICE uses the binary concrete vectors to model the variance of the effect sizes. This property allows our model to adjust for infinitesimal effects from the non-causal variants. Figure 10 compares the average run-time of each method across all parameter settings. We observe that the run-time of BEATRICE and SuSiE are less than one minute. FINEMAP requires five minutes on average to converge. CARMA requires approximately 100 minutes to converge, likely due to the slow MCMC sampling procedure for generating posterior estimates.

**Fig. 10.**
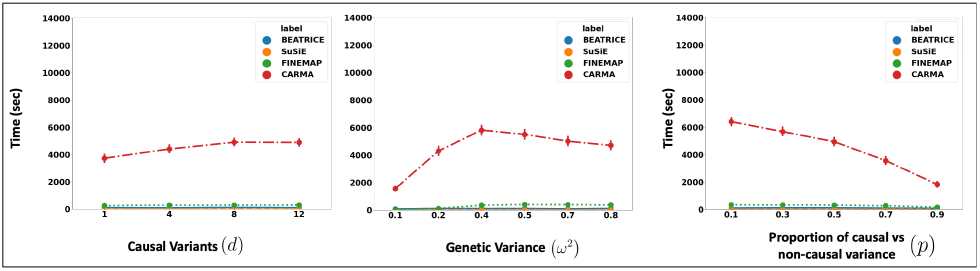
The runtime comparison of BEATRICE, SuSiE, and FINEMAP across all the simulation settings.

The final contribution of BEATRICE is its simple and flexible design. Importantly, BEATRICE can easily incorporate priors based on the functional annotations of the variants. Specifically, in the current setup, the prior over **c** is effectively constant, as captured by 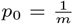. We can integrate functional information simply by modifying the distribution of *p*_0_ across the variants. Going one step further, a recent direction in fine-mapping is to aggregate data across multiple studies to identify causal variants [20]. This extension would amount to adding multiple log-likelihood terms in Eq. (12) corresponding to the different GWAS inputs 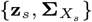 matrices across studies. A similar extension can be used for multiple ancestries or traits.

In this case, the inputs 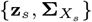 would reflect a particular ancestry or trait. The main difference between BEATRICE and these extensions is that the parameters **p** would be generated as a function of multiple **z**_*s*_ scores, i.e., **p** = *f* (**z**_1_, …, **z**_*S*_; *ϕ*).

In this work, we have shown that BEATRICE is highly efficient in handling the complexity that arises due to infinitesimal effects and out-of-sample LD matrices (**Section S1.13**). Thus, we believe that the advantages of BEATRICE will be more evident when considering polygenic traits and diseases. Additionally, the high coverage and small size of credible sets reported in Figure 2–4 show that BEATRICE can successfully prioritize variants in the presence of LD. This property is in stark contrast with the baseline finemapping approaches that generate a large number of credible sets that do not contain a causal variant. Taken together, we believe BEATRICE could be useful in eQTL studies, where multiple variants within a locus can show strong association due to the complex LD structure present in the human genome [39].

In summary, we present BEATRICE, a novel Bayesian framework for fine-mapping that identifies potentially causal variants within GWAS risk loci through the shared LD structure. Using a variational approach, we approximate the posterior probability of the causal location(s) via a binary concrete distribution. In conjunction, we introduce a new strategy to build a reduced set of causal configurations within the exponential search space that can be neatly folded into our optimization routine. This reduced set is used to approximate the PIPs and identify credible sets. We have demonstrated through a comprehensive simulation the advantages of BEATRICE under different noise settings and that BEATRICE outperforms existing fine-mapping methods. Hence, BEATRICE is a powerful tool to refine the results of a GWAS or eQTL analysis. It is also flexible enough to accommodate a variety of experimental settings.

## Acknowledgments

This work was supported by the National Science Foundation CAREER Award 1845430 (PI: Venkataraman), the National Institutes of Health Awards R01-HD108790 (PI: Venkataraman), U24-HG010263 (PI: Schatz) and U41-HG006620 (PI: Schatz).

## Code Availability

We have compiled the code for BEATRICE and its dependencies into a docker image, which can be found at https://github.com/sayangsep/Beatrice-Finemapping. We have also provided details about the usage and outputs of the model in Supplementary **Section S1.20**.

## Authors Declaration

The authors do not have any conflicting interests.

## Footnotes

1 https://github.com/sayangsep/Beatrice-Finemapping

2 https://registry.opendata.aws/ukbb-ld/

