## Supplementary Methods for "BEATRICE: Bayesian Fine-mapping from Summary Data using Deep Variational Inference"


### S1 Detailed Methods and Additional Applications

#### S1.1 Distribution of Normalized GWAS Effect Sizes

GWAS uses a collection of element-wise linear regression models to estimate the effect of each genetic variant. Mathematically, the collection of GWAS effect sizes are computed as $\hat{\beta} = \frac{1}{n}\mathbf{X}^T\mathbf{y}$, with the corresponding vector of normalized z-scores equal to $\mathbf{z} = \sqrt{\frac{\tau}{n}}\mathbf{X}^T\mathbf{y}$ [1, 2]. Under additive linear assumption of the quantitative trait (Section 2.1), we can write the normalized effect sizes as follows:

$$\mathbf{z} = \sqrt{\frac{\tau}{n}}\mathbf{X}^T\left(\mathbf{X}\beta + \eta\right) \text{ where } \eta \sim N\left(0, \frac{1}{\tau}\mathbf{I}_n\right) \tag{1}$$

$$\Longrightarrow P\left(\mathbf{z}|\beta\right) = N\left(\mathbf{z}; \sqrt{(n\tau)}\beta\mathbf{\Sigma}_X, \mathbf{\Sigma}_X\right) \tag{2}$$

Under the assumptions of our generative process, the prior on the effect sizes $\beta$ follows a normal distribution $N\left(0, \frac{1}{\tau}\sigma^2\mathbf{\Sigma}_C\right)$. Under this assumed prior, the posterior distribution of the z-scores $\mathbf{z}$ can be written as follows:

$$p\left(z|\mathbf{\Sigma}_X, \mathbf{\Sigma}_C\right) = N\left(\mathbf{z}; 0, \mathbf{\Sigma}_X + \mathbf{\Sigma}_X\left(n\sigma^2\mathbf{\Sigma}_C\right)\mathbf{\Sigma}_X\right) \tag{3}$$

#### S1.2 Properties of Binary Concrete Random Vectors

Each element of a binary concrete random vector can be viewed as a continuous relaxation of a Bernoulli random variable that can be mathematically represented as

$$\mathbf{c}_i = \xi\left(\frac{\log\left(\frac{U}{1-U}\right) + \log\left(\frac{\mathbf{p}_i}{1-\mathbf{p}_i}\right)}{\lambda}\right), \tag{4}$$

where $\xi(\cdot)$ is a sigmoid function, $\lambda$ controls the extent of relaxation from Bernoulli random variable, $U$ is a uniform random variable that introduces randomness, and $\mathbf{p}_i$ is an approximate probability measure of finding a causal variant at location $i$. As shown in Figure S1(a) smaller values of $\lambda$ lead to progressively discretized $\mathbf{c}_i$, while larger values provide a smoother mapping. This continuous representation allows us to model the infinitesimal effects of the non-causal variants. In Figure S1(b) we show how the

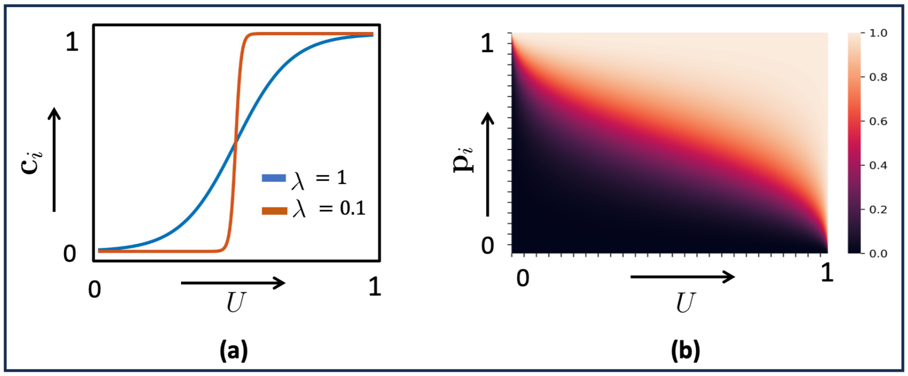


**Fig S1.** Properties of the binary concrete distribution. (a) Relationship between $\mathbf{c}_i$ and $U$ for different values of $\lambda$. (b) The change in $\mathbf{c}_i$ for varying probability map value $\mathbf{p}_i$ and uniform noise $U$. The darker and brighter colors represents $\mathbf{c}_i$ close to 0 and 1, respectively.

joint relationship between $\mathbf{p}_i$ and the sampled uniform random variable $U$ generates the binary concrete random variable $\mathbf{c}_i$. As seen, higher values of $\mathbf{p}_i$ push $\mathbf{c}_i$ closer to 1, irrespective of the uniform random variable. Intuitively, a variant with high probability is a likely candidate to contain a causal variant.

#### S1.3 Identification of Credible Sets for BEATRICE

One of the notable features of BEATRICE is its ability to identify a comprehensive set of causal configurations with non-negligible posterior probability within the exponentially large search space. As described in Section 3.1, the reduced search space $\mathcal{B}^R$ is comprised of vectors that BEATRICE randomly samples at each iteration of the optimization. We identify credible sets from $\mathcal{B}^R$ in two steps.

First, in a sequential fashion, we identify the "key" variant with the highest conditional probability given the previously selected variants. Formally, let $\mathcal{K}$ be the indices of previously identified "key" variants. The conditional probability for variant $i$ given $\mathcal{K}$ in each iteration can be calculated as follows:

$$P\left(\mathbf{b}_i = 1 | \mathcal{K}, \mathbf{z}, \mathbf{\Sigma}_X\right) = \frac{\sum_{\mathbf{b} \in \mathcal{C}} P\left(\mathbf{b} | \mathbf{z}, \mathbf{\Sigma}_X\right)}{\sum_{\mathbf{b}' \in \mathcal{D}} P\left(\mathbf{b}' | \mathbf{z}, \mathbf{\Sigma}_X\right)} \tag{5}$$

$$\begin{aligned} \text{s.t. } & \mathcal{D} \subset \mathcal{B}^R \text{ and } \mathcal{D} = \{\mathbf{b} | \mathbf{b}_j = 1 \ \forall \ j \in \mathcal{K}\} \\ & \mathcal{C} \subset \mathcal{B}^R \text{ and } \mathcal{C} = \{\mathbf{b} | \mathbf{b}_j = 1 \ \forall \ j \in \{i\} \cup \mathcal{K}\} \end{aligned}$$

where, $\mathcal{D}$ is the subset of $\mathcal{B}^R$ that includes all of "key" variants and $\mathcal{C}$ is the subset of $\mathcal{B}^R$ that includes both variant $i$ and the "key" variants.

We perform this sequential variant selection until the maximum posterior probability reduces below a threshold, which we define as the "key" threshold $\gamma_{key}$ and fix at $\gamma_{key} = 0.2$ for all experiments. We note that this threshold can be controlled by the user. The selected "key" variants act as proxy for highly plausible causal variants.

In the second step, we identify the set of variants that can replace the "key" variant in the causal configurations while maintaining a high posterior probability. This set of variants act as a credible set for that particular "key" variant. To do this, we first remove one of the key variants from $\mathcal{K}$ and estimate the posterior probability of other variants given the remaining "key" variants. For example, let variant $k_1$ be a " key"

**Algorithm 1** Algorithm to find credible sets

$\mathcal{K} = \{\}$
$\mathcal{CS} = \{\}$
Estimate posterior probabilities accoding to Eq. 5.
**while** $max\left[P(\mathbf{b}_i|\mathcal{K}, \mathbf{z}, \mathbf{\Sigma}_X) \mid \forall i \notin \mathcal{K}\right] > \gamma_{key}$ **do**
    $\mathcal{K} = \mathcal{K} \cup argmax\left[P(\mathbf{b}_i|\mathcal{K}, \mathbf{z}, \mathbf{\Sigma}_{\mathbf{X}}) \mid \forall i \notin \mathcal{K}\right]$
    Estimate posterior probabilities accoding to Eq. 5.
**end while**
**for** $k \in \mathcal{K}$ **do**
    $\mathcal{S} = \{\}$
    $cov = 0$
    Generate $\mathcal{K}'$ by removing $k$ from "key" set.
    **for** $i = [1, \ldots, m]$ and $i \notin \mathcal{K}'$ **do**
        **if** allow_dup == False **then**
            Remove $i$-th SNPs $\forall i \in \mathcal{CS}$
        **end if**
        Estimate posterior probability according to Eq. 6
        Stack the probability scores in a vector $\mathcal{P}$.
    **end for**
    **while** $max\left[\mathcal{P}_i \mid \forall i \notin \mathcal{K}'\right] > \gamma_{selection}$ and $cov < \gamma_{coverage}$ **do**
        $\mathcal{S} = \mathcal{S} \cup argmax\left[\mathcal{P}_i \mid \forall i \notin \mathcal{K}'\right]$
        $cov = cov + max\left[\mathcal{P}_i \mid \forall i \notin \mathcal{K}'\right]$
    **end while**
    Add $\mathcal{S}$ to $\mathcal{CS}$ as credible set of $k$.
**end for**

variant. We estimate the posterior probabilities as follows:

$$P\left(\mathbf{b}_i = 1|\mathcal{K}', \mathbf{z}, \mathbf{\Sigma}_X\right) = \frac{\sum_{\mathbf{b}\in\mathcal{G}} P\left(\mathbf{b}|\mathbf{z}, \mathbf{\Sigma}_X\right)}{\sum_{\mathbf{b}'\in\mathcal{H}} P\left(\mathbf{b}'|\mathbf{z}, \mathbf{\Sigma}_X\right)} \tag{6}$$

$$\begin{aligned} \text{s.t. } & \mathcal{K} = \mathcal{K}' \cup \{k_1\} \\ & \mathcal{G} = \{\mathbf{b}|\mathbf{b}_j = 1 \ \forall\ j \in \{i\} \cup \mathcal{K}'\} \\ \mathcal{H} \subset \mathcal{B}^R \text{ and } & \mathcal{H} = \{\mathbf{b}|\mathbf{b}_j = 1 \ \forall\ j \in \mathcal{K}'\} \end{aligned}$$

where $\mathcal{K}'$ is the set of "key" variants without $k_1$, $\mathcal{G}$ is the set of configurations that include both variant $i$ and the remaining "key" variants, and and $\mathcal{H}$ is the set of causal configurations that include all "key" variants except $k_1$.

Once computed, we sort these posterior probabilities in descending order and add the corresponding variants to the credible set until the cumulative sum reaches the coverage threshold $\gamma_{coverage}$. By default, we do not allow overlap between the credible sets. However, this setting can be relaxed using the flag *allow_dup* when calling BEATRICE. We fix the coverage threshold at $\gamma_{key} = 0.95$ in this work, but it too can be set by the user. Finally, we prune uncorrelated variants by thresholding the posterior probability according to the selection threshold $\gamma_{selection} = 0.05$, again a tunable parameter for users. Algorithm 1 provides a detailed description of these steps.

#### S1.4 Detailed Architecture of Inference Module

BEATRICE consists of three main components: an inference module, a random sampler, and a generative module. Figure S2 shows the detailed neural network architecture of

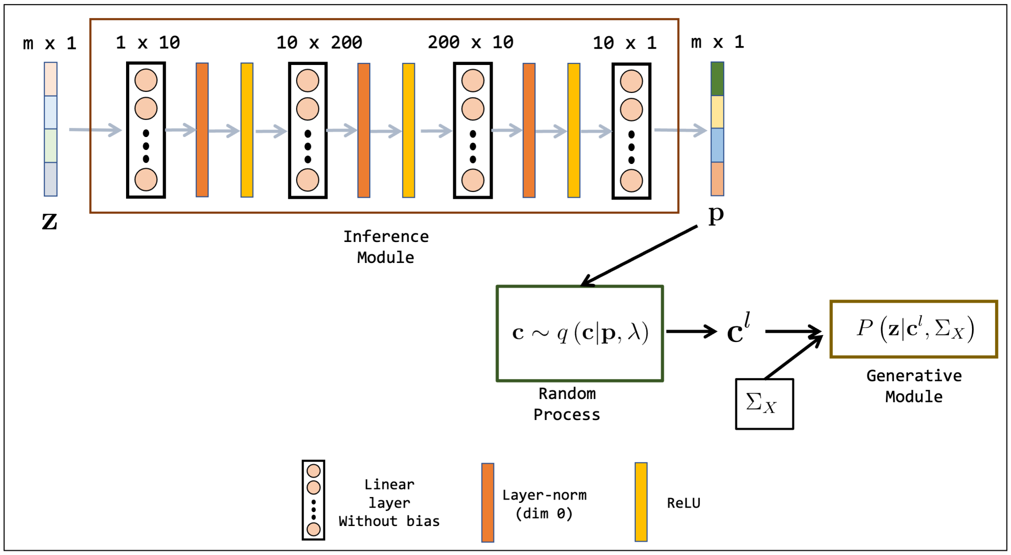


**Fig S2.** Neural network architecture for the inference module used in BEATRICE. The neural network uses a sequence of linear layers, layer normalization, and activation layers. The dimension of the linear layers are shown on top of each layer. The input to the inference module is the normalized z-scores obtained from GWAS. The output of the inference module is the estimated parameters of our binary concrete distribution.

the inference module. The neural network is trained to output the parameters $\mathbf{p}$ of the binary concrete distribution that we use to approximate the posterior distribution of the causal configurations given the z-statistics and LD matrix. We use the variable $\phi$ to denote the collection of learnable weights in the neural network. The weights $\phi$ are trained using gradient descent to minimize the KL divergence loss given in Eq. (9) of the manuscript. This process goes as follows: given an an input z-statistic (1) the neural network generates parameters $\mathbf{p}(\theta)$; (2) the random process sampler uses these parameters to generate the causal configuration $\mathbf{c}$ according to Eq. (6); (3) the generative module uses Eq. (4) to compute the log-likelihood. Finally, we compute the KL divergence loss and use gradient descent to update the neural network weights $\phi$. This three-step process is repeated until convergence. Notice that $\phi$ is a function of $\mathbf{z}$ and $\mathbf{\Sigma}_X$ because the KL divergence loss is a function of both these inputs. This property ensures that the neural network uses the data to generate optimal parameters $\mathbf{p}$ for our proposal distribution $q(\cdot; \mathbf{p}, \lambda)$. The implicit function learned by this network helps BEATRICE to handle complex and possibly nonlinear interactions in the data without being constrained by parametric representations.

#### S1.5 Computational Complexity

Each iteration of stochastic gradient descent requires us to compute the data log-likelihood term $\left[\log N\left(\mathbf{z}; 0, \mathbf{\Sigma}_X + \mathbf{\Sigma}_X\left(n\,\sigma^2\,\mathbf{\Sigma}_C^l(\phi)\right)\mathbf{\Sigma}_X\right)\right]$. This computation is expensive due to the covariance matrix inversion, whose run-time is on the order of $O(m^3)$, where $m$ is the total number of variants. To mitigate this issue, the works of [3] show that if $\mathbf{\Sigma}_C^l(\phi)$ is sparse, then the matrix inversion can be done with order $O(k^3) + O(mk^2)$ run-time, where $k$ is the number of non-zero diagonal elements of $\mathbf{\Sigma}_C^l(\phi)$. We leverage this result in the optimization by thresholding the elements of $\mathbf{c}^l(\phi)$ to set small values exactly to zero. In every iteration, we sparsify $\mathbf{c}_t^l$ by considering the top 50 non-zero locations of $\mathbf{c}_t^l$

with values $\mathbf{c}_t^l(i) > 0.01$. This strategy provides a way to optimize the parameters of our models in $O(50^3) + O(m50^2)$ run-time for all scenarios. We also regularize $\mathbf{\Sigma}_X$ with a small diagonal load to ensure invertibility of the covarance matrix at each iteration. Finally, we run stochastic gradient descent with a batch size of one to further speed up BEATRICE. Effectively, this means that we sample a single $\mathbf{c}^l(\phi)$ at each epoch rather than perform a true Monte Carlo integration. The authors of [4] have previously shown that a single random sample ($L = 1$) is sufficient to guarantee convergence to a local minimum of Eq. (12) reported in the main text. Algorithm **??** in the main text provides a detailed description of these optimization steps.

#### S1.6 Finemapping Under Varying Phenotypic Variance Explained by Non-causal SNPs

In this section we probe finemapping under varying SNP heritability captured by two simulation settings: $\{d = 1, \omega^2 = 0.1, p = 0.3\}$ (Figure S3) and $\{d = 1, \omega^2 = 0.4, p = 0.1\}$ (Figure S4). In the first case, non-causal SNps explain 7% of the observed phenotypic variance, and in the second case, they explain 36% of the phenotypic variance. Under both settings, the causal SNP has a lower z-score, as compared to the neighboring variants. Figure S3 describes the first setting. In this case, the variance explained by the non-causal variants is small, making it easy for all three methods to correctly identify the true casual SNP and assign it the highest PIP. Figure S4 describes the second setting. Here, the non-causal SNPs have much higher effect sizes than the true causal SNP. Correspondingly, we observe that only BEATRICE is the only method that assigns the highest PIP to the true causal SNP. Both FINEMAP and SuSiE generate uncertain predictions, as captured by the large, credible sets and multiple high PIPs. The high z-scores observed for the non-causal SNPs in Figure S3 and Figure S4 can be largely attributed to the LD structure present between SNPs. Following the generative assumptions of fine-mapping in Eq. (3), we can show that the estimated effect size $\hat{\beta}_i$ for a given variant $i$ can be expressed as $\hat{\beta}_i = \sum_j r_{ij}\beta_j$, where $r_{ij}$ is the correlation between variants $i$ and $j$, and $\beta_j$ is the true effect size. This expression reinforces that in the presence of an infinitesimal effect (i.e., $\beta_j \neq 0$), the LD structure can inflate the estimated size of a variant, leading to a high z-score. We conjecture that BEATRICE uses the binary concrete distribution to model non-causal variants with non-zero effects, while using the sparsity term of $\mathcal{L}(\cdot)$ to prioritize potentially causal variants.

#### S1.7 Identification of Credible Sets for FINEMAP

FINEMAP outputs a collection of credible sets under the assumption of multiple causal variants $d = 1, \ldots, D$. Similar to the approach used in [5], we sub-select the credible sets from this collection with the highest posterior probability. From here, we pruned the sets with minimum absolute purity greater that 0.5. As defined in [5], purity is the pairwise correlation coefficient between the variants, obtained from the LD matrix.

#### S1.8 FDR vs. Power Curves for Different Values of SNP Heritability ($\omega^2$)

Figure S5–S7 show the FDR vs. power performance across models for SNP heritability $\omega^2 = 0.2$, $\omega^2 = 0.4$ and $\omega^2 = 0.8$, respectively. Here we show the mean performance of all the methods across 20 simulation settings with fixed SNP heritabilities and multiple causal variants $d = \{1, 4, 8, 12\}$ and multiple infinitesimal effects from non-causal SNPs $p = \{0.1, 0., 3, 0.5, 0.9\}$. We use linear interpolation to project the results across

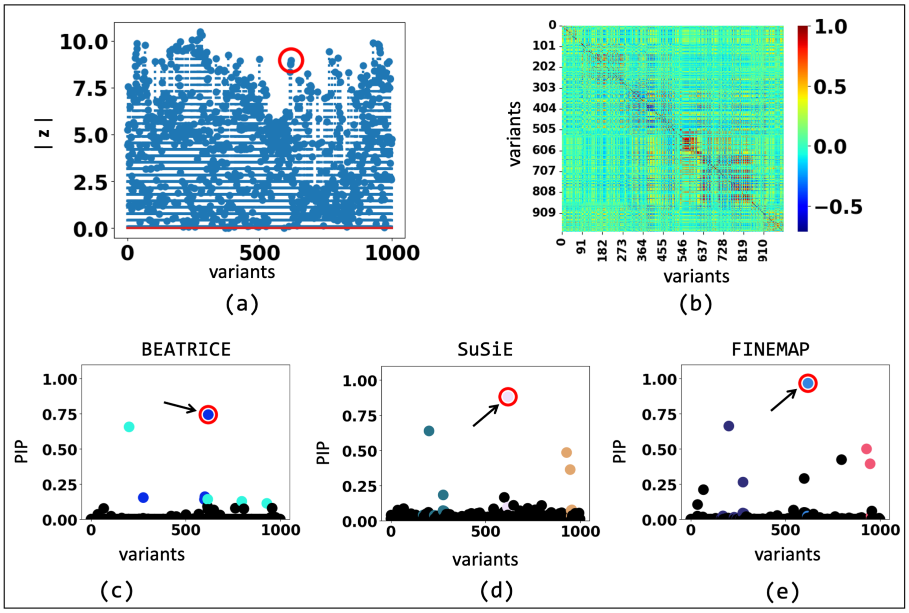


**Fig S3.** The fine-mapping performance of BEATRICE , SuSiE, and FINEMAP at a noise setting of $\{d = 1, \omega^2 = 0.1, p = 0.3\}$. (a) The absolute z-score of each variant as obtained from GWAS. (b) Pairwise correlation between the variants. (c)-(e) illustrate the posterior inclusion probabilities of each variant, as estimated by the three method. The red circle marked by an arrow shows the location of the causal variant. The non-black markers represent the variants assigned to a credible set, color-coded based on the assignment.

simulation runs onto the same x-y axes. When $p$ is small, i.e., the scenario in which most of the phenotype variance can be explained by the infinitesimal effects from non-causal variants, we notice that BEATRICE gives the best performance in terms of power and FDR. This result shows the BEATRICE generates PIPs that are robust in the presence of high infinitesimal effects from nearby SNPs. This property shows BEATRICE is consistent across different SNP heritability values. These results suggest that BEATRICE can better estimate the causal variant(s) in the presence of confounding information from non-causal variants.

#### S1.9 Detailed Comparison Analyses

In this section, we compare the performances of the models across individual noise settings and provide further insight about the advantages of BEATRICE. Figure S9, Figure S10, and Figure S8 show the performance comparison of AUPRC, power and coverage, respectively. Figure S8 shows a significant improvement in coverage compared to the baselines across noise settings. In addition, the BEATRICE shows uniformly better AUPRC in Figure S9. However, in terms of power, all models exhibit similar performance. A high coverage with comparable power suggests that BEATRICE can identify high quality credible sets that contain causal variants. In contrast, the baselines identify many credible that do not contain a causal variant, ultimately leading to low coverage.

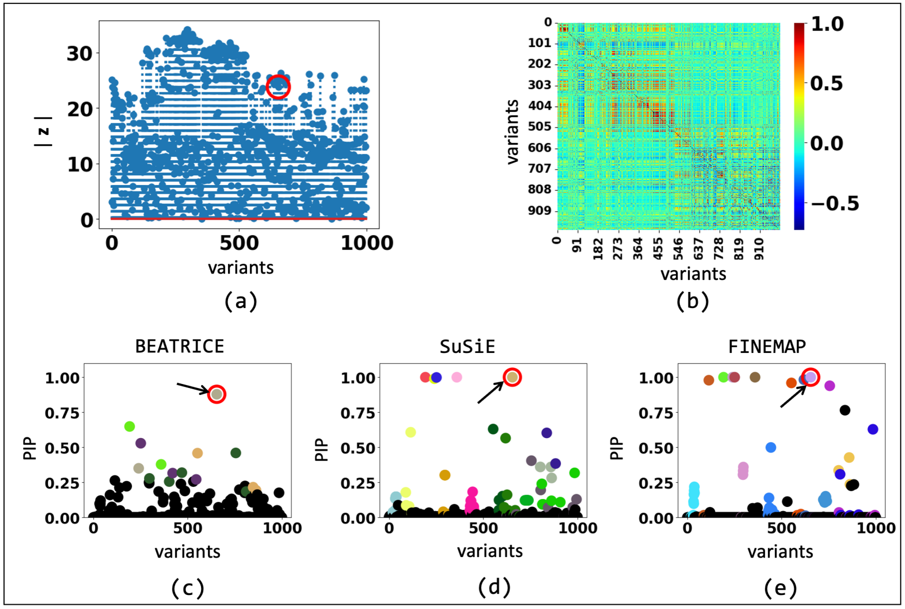


**Fig S4.** The fine-mapping performance of BEATRICE , SuSiE, and FINEMAP at a noise setting of $\{d = 1, \omega^2 = 0.4, p = 0.1\}$. (a) The absolute z-score of each variant as obtained from GWAS. (b) Pairwise correlation between the variants. (c)-(e) illustrate the posterior inclusion probabilities of each variant, as estimated by the three method. The red circle marked by an arrow shows the location of the causal variant. We have further color-coded the variants based on their assignment to credible sets. The non-black markers represent the variants assigned to a credible set, color-coded based on the assignment.

#### S1.10 Performance Across Multiple Thresholds of $\gamma$

Supplementary Figures S11–S13 show the performance of BEATRICE for three different values of the hyperparameter $\gamma$, namely, $\gamma = \{0.01, 0.1, 0.5\}$. We observe that a smaller value of $\gamma$ leads to larger credible sets, which is expected because of the increases in the number of probable causal configurations. We also observe that a decrease in the value of $\gamma$ leads to increased power and reduced coverage. This occurs a lower threshold results in more SNPs identified as causal. This scenario produces low-quality credible sets, as many of them will not contain any causal SNP (i.e., low coverage). On the other hand, since more SNPs are identified as causal, we are more likely to select the ground-truth causal signal (i.e., high power).

#### S1.11 Calibration of PIPs

In this section, we study the calibration of the PIPs generated by the three fine-mapping approaches. Calibration is defined as the proportion of causal SNPs that lie within a bin of SNPs with a fixed PIP. Figure S14 illustrates the calibration of SNPs that lie within five different PIP bins, as aggregated across our 2400 simulation experiments. Figure S14 shows a similar trend of miscalibration in the presence of infinitesimal from non-causal SNPs reported in the recent work of [6]. However, we observe that BEATRICE shows significantly better calibration compared to the other approaches. This result highlights that BEATRICE can successfully account for multiple causal SNPs in the presence of infinitesimal effects for other variants.

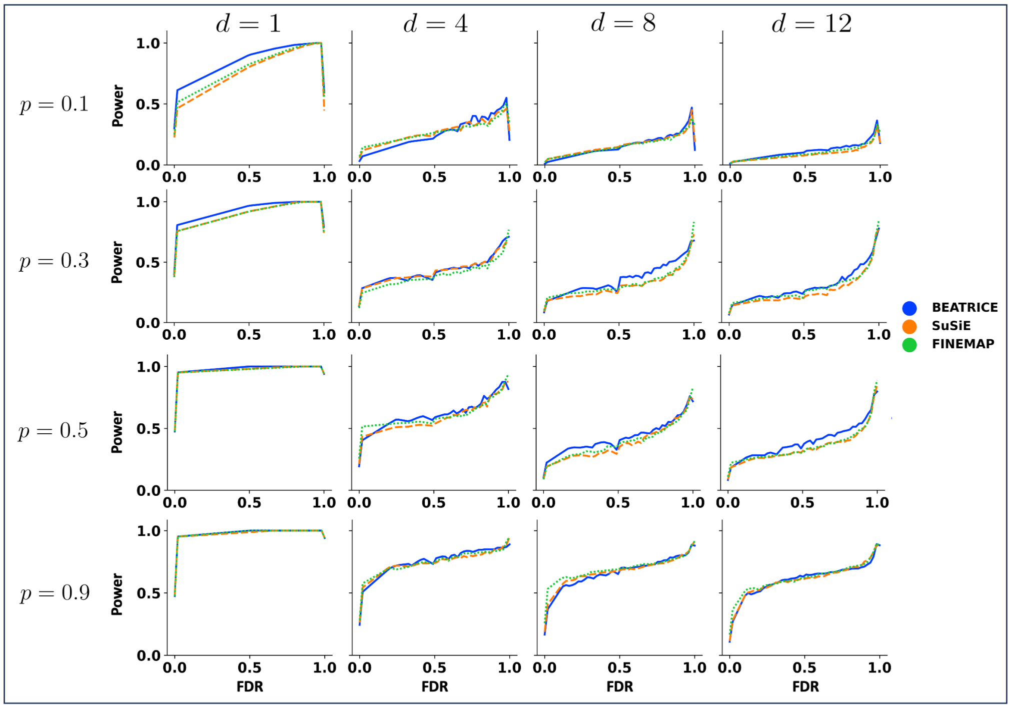


**Fig S5.** Power vs. FDR curve for three models across multiple causal variants $d = [1, 4, 8, 12]$, and multiple proportion of phenotype variance explained by causal variants $p = [0.1, 0.3, 0.5, 0.9]$, while fixing SNP heritability at $\omega^2 = 0.2$. Each row and column corresponds to a specific value of $p$ and $d$, respectively. In each plot, the y-axis captures power, and the x-axis represents FDR.

#### S1.12 Credible Curves Across Constant Power

Fig. S15 reports the change in AUPRC, coverage, and size of credible sets for constant power across 2400 simulation experiments. The results highlight that with increasing power, the size of credible sets obtained from BEATRICE remains smaller or comparable to all the other baselines. Moreover, compared to SuSiE and FINEMAP, we see a significant improvement in coverage and AUPRC for BEATRICE. These experimental results validate that BEATRICE generates better credible sets and PIPs that lead to improved coverage and AUPRC, respectively.

#### S1.13 Finemapping In The Presence of Out-of-sample LD Matrices

Figure S16 shows the performances of BEATRICE , SuSiE, and FINEMAP with both in-sample and out-of-sample LD matrices across multiple causal variants. In this experiment, we first generate $10,000$ genetics samples according to the procedure described in Section 4.1. We then use $5,000$ samples to generate the $\mathbf{z}$ score and phenotype. The remaining $5,000$ samples are used to generate the out-of-sample LD matrices. While we observe a drop in performance across all methods when using an out-of-sample LD matrix, this drop is more significant for SuSiE and FINEMAP. Specifically, when using an in-sample LD matrix, the AUPRC of SuSiE is similar to that of BEATRICE. However, the performance of SuSiE degrades substantially when using an out-of-sample LD matrix, even when compared to the reduced performance of BEATRICE. We also observe that BEATRICE yields the best coverage with similar

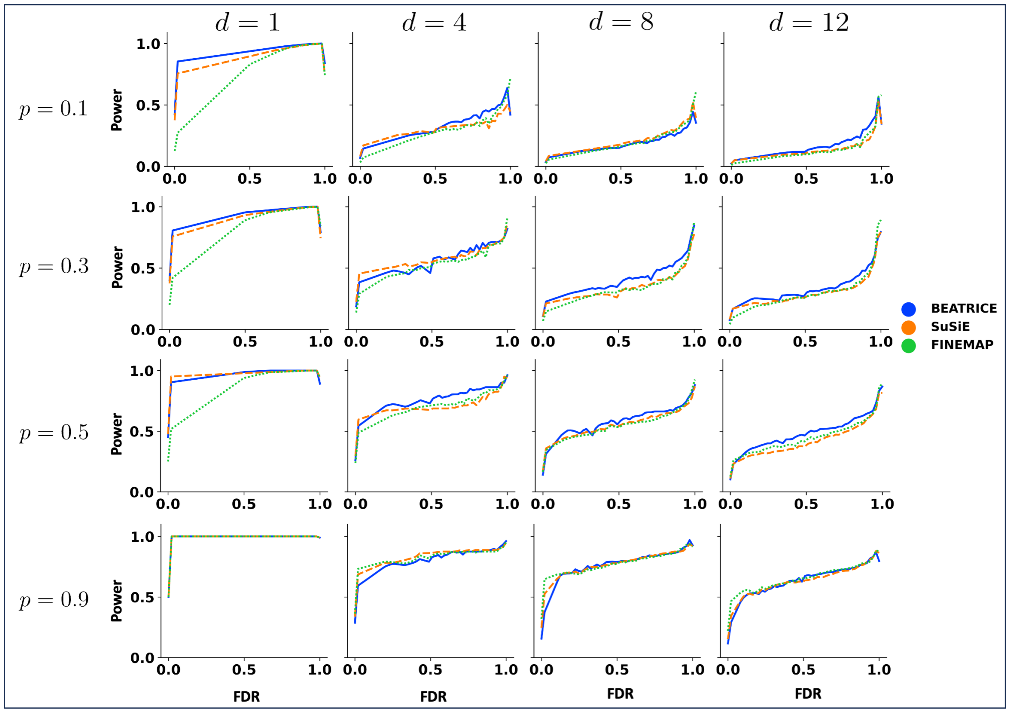


**Fig S6.** Power vs. FDR curve for three models across multiple causal variants $d = [1, 4, 8, 12]$, and multiple proportion of phenotype variance explained by causal variants $p = [0.1, 0.3, 0.5, 0.9]$, while fixing SNP heritability at $\omega^2 = 0.4$. Each row and column corresponds to a specific value of $p$ and $d$, respectively. In each plot, the y-axis captures power, and the x-axis represents FDR.

power compared to SuSiE and FINEMAP. This result demonstrate that BEATRICE can estimate more accurate PIPs and credible sets, leading to improved performance, even in the presence of LD mis-specification.

#### S1.14 Evaluating the Impact of Covariates

The summary statistics in GWAS are often calculated after adjusting for multiple covariates like age, sex, and genetic PCS. Accordingly, we use simulated data to compare the effect of these covariates on the finemapping approaches.

We note that a GWAS solves the following mathematical relationship:

$$\mathbf{y} = \mathbf{x}_i\beta_i + \mathbf{C}\gamma + \epsilon \tag{7}$$

where $\mathbf{x}_i$ is the genetics information across samples for the $i$-th SNP, $\beta_i$ is the GWAS effect size for the $i$-th SNP, $\mathbf{C}$ is the covariate matrix, and $\gamma$ are the covariate regression coefficients.

The model in Eq. (7) can be simplified by eliminating the covariates using the projection matrix $\mathbf{P} = \mathbf{I} - \mathbf{C}(\mathbf{C}^T\mathbf{C})^{-1}\mathbf{C}^T$ [7]. Multiplying Eq. (7) with $\mathbf{P}$ yields the following:

$$\widetilde{\mathbf{y}} = \widetilde{\mathbf{x}}_i\beta_i + \epsilon^* \tag{8}$$

where $\widetilde{\mathbf{x}}_i = \mathbf{P}\mathbf{x}$ and $\widetilde{\mathbf{y}} = \mathbf{P}\mathbf{y}$ are the covariate-adjusted genetics data and phenotype, respectively. Eq. (8) exactly matches our fine-mapping framework. Thus, computing the LD matrix from the original genetics data instead of the covariate-adjusted data amounts to an LD matrix mis-specification related to the projection matrix $\mathbf{P}$.

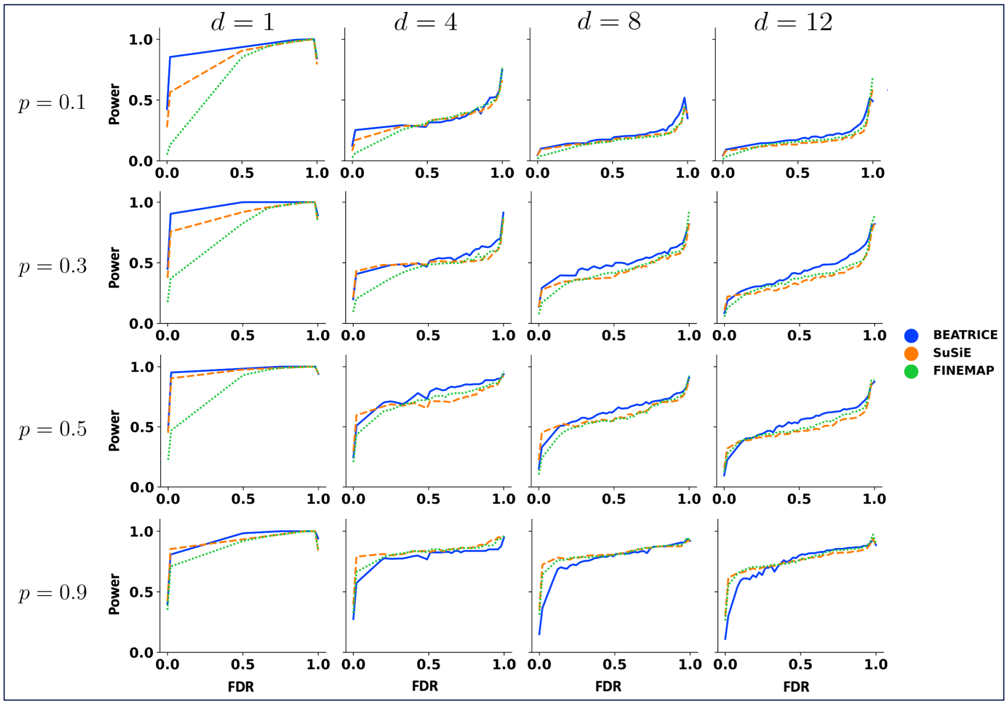


**Fig S7.** Power vs. FDR curve for three models across multiple causal variants $d = [1, 4, 8, 12]$, and multiple proportion of phenotype variance explained by causal variants $p = [0.1, 0.3, 0.5, 0.9]$, while fixing SNP heritability at $\omega^2 = 0.8$. Each row and column corresponds to a specific value of $p$ and $d$, respectively. In each plot, the y-axis captures power, and the x-axis represents FDR.

To study the effect of this type of LD misspecification, we generate the phenotypic data according to Eq. (7), where the covariate matrix $\mathbf{C}$ is taken as the 10 principal components (PCs) obtained from the genetic data. Principal components are often used to adjust for population-level effects in genetic studies [8]. Following the strategy described in Section 4.1 we fix the phenotypic variance explained by genetics at 0.5 and the phenotypic variance explained by covariates at 0.2. We then sweep over the number of causal variants and the ratios of phenotypic variance explained by causal and non-causal SNPs. The results of this experiment are shown in Fig. S17. We observe the same performance trends as in our main simulated experiments, i.e., BEATRICE obtains the best AUPRC and coverage with comparable power.

With regards to real-world experiments, the UK biobank dataset directly adjusts for the covariates and generates LD matrices, which are publicly available [9] and helps to mitigate the mis-specification. Still, our real-world experiments suggest that BEATRICE can find potential causal variants (e.g., apoe-$\epsilon$2) for AD in given external covariates and out-of-sample LD matrix.

#### S1.15 Performance Comparison When Allowing Overlap in the Credible Sets

In this Section, we compare the performance of BEATRICE , FINEMAP, and SuSiE when allowing for overlap between credible sets. In the case of BEATRICE , we enable this setting through a flag in the function call. In contrast, SuSiE and FINEMAP do not have an explicit flag to control the credible sets. Nonetheless, we notice empirically that among 2400 simulation experiments, there are 21 and 29 scenarios where

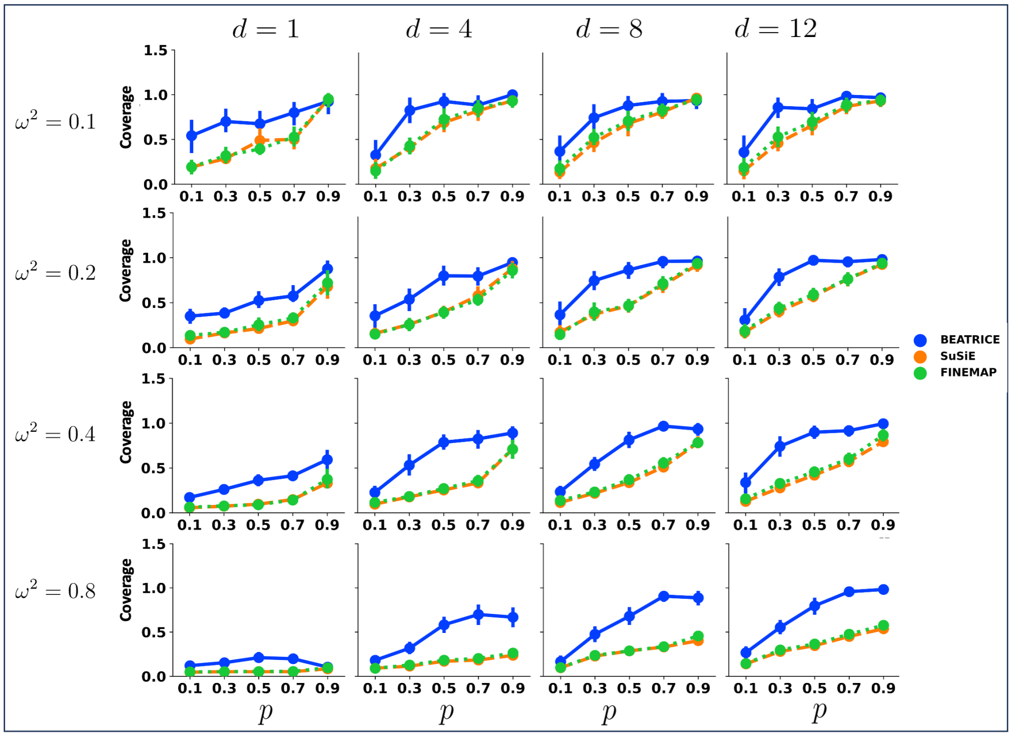


**Fig S8.** Coverage of the credible sets generated by the three models across multiple causal variants $d = [1, 4, 8, 12]$, multiple SNP heritability $\omega^2 = [0.1, 0.2, 0.4, 0.8]$ and multiple infinitesimal effects from non-causal variants $p = [0.1, 0.3, 0.5, 0.7, 0.9]$. Each row and column corresponds to a specific value of $\omega^2$ and $d$, respectively. In each plot, the y-axis captures coverage, and the x-axis represents $p$.

FINEMAP, and SuSiE, have overlapping variants across credible sets. Such scenarios can arise when two credible sets, estimated for different causal variants, contain other variants with high LD. Similar to Section 4.3.1, we report the AUPRC, coverage, power, and size while sweeping over the number of causal variants $d$, the genotype variance $\omega^2$, and the percentage of phenotypic variance explained by the causal SNPs $p$.

Figure S18, Figure S19, and Figure S20 report the performance of each model for different values of $d$, $\omega^2$, and $p$, respectively. As seen, BEATRICE has slightly lower power than when enforcing non-overlapping credible sets (main text), but it remains within the 95% confidence interval of both SuSiE and FINEMAP. This slight difference occurs because SuSiE and FINEMAP generate a larger number of credible sets as compared to BEATRICE, with many of them not containing a causal variant. This scenario allows SuSiE and FINEMAP to cover many variants, improving power. At the same time, the coverage of these methods is much lower. The trends in AUPRC and credible set size are similar to the case of non-overlapping sets.

#### S1.16 Comparison with SuSiE-inf

SuSiE-inf [10] is an extension of the SuSiE model that accounts for infinitesimal effects from non-causal variants. In this section, we compare the performance of BEATRICE with SuSiE-inf across the same simulation setting as in Section 4.1 of the main text.

Unlike BEATRICE, we observe that SuSiE-inf fails to converge in multiple cases. Specifically, Figure S21 illustrates the number of experimental settings, for which SuSie-inf fails to converge across each parameter sweep. This problem becomes

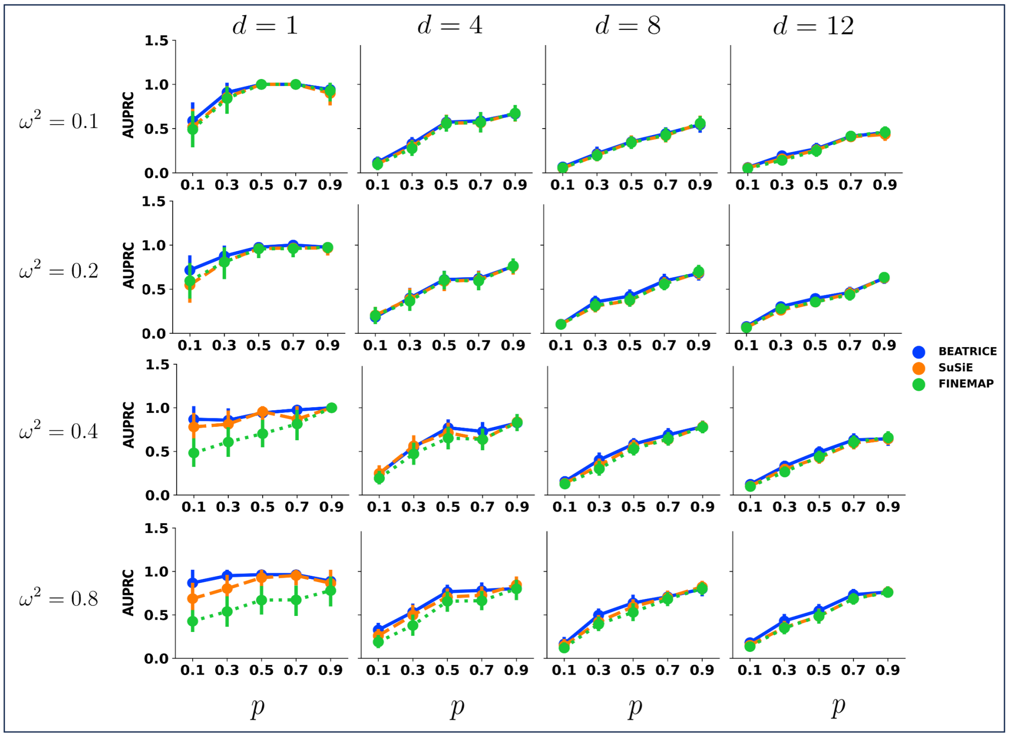


**Fig S9.** AUPRC of PIPs generated by the three models across multiple causal variants $d = [1, 4, 8, 12]$, multiple SNP heritability $\omega^2 = [0.1, 0.2, 0.4, 0.8]$ and multiple infinitesimal effects from non-causal variants $p = [0.1, 0.3, 0.5, 0.7, 0.9]$. Each row and column corresponds to a specific value of $\omega^2$ and $d$, respectively. In each plot, the y-axis captures AUPRC, and the x-axis represents $p$.

prominent with increasing SNP heritability, as explained by $\omega^2$.

Figure S22, Figure S23, and Figure S24 shows the performance comparison between BEATRICE and SuSiE-inf. We emphasize that the performance of SuSiE-inf is computed based only on the convergent runs, so these values should be treated as optimistic. In contrast, the performance of BEATRICE is computed across all runs, as we did not face convergence issues with our model. Across different parameter sweeps, we see that the coverage of SuSiE-inf is similar to BEATRICE. However, BEATRICE achieves uniformly better power and AUPRC.

#### S1.17 Functional Annotation of Finemapped SNPs Obtained from The Alzheimer's Study

In an exploratory analysis, we investigate the biological consequences of the SNPs with high PIPs ($> 0.9$) of the first clump, as identified by each method. Details about the SNPs identified by BEATRICE and SuSiE are provided in Supplementary Table 2 and Supplementary Table 3, respectively. We also provide the p-values from the GWAS statistics and the PIPs identified by each method. We extracted all genes tagged by BEATRICE and SuSiE from the Ensemble VEP annotation, which expands the GENCODE boundaries by 5kb to account for upstream/downstream flanking regulatory regions. Supplementary Table 4 and Supplementary Table 5 show the tagged genes, and the biological consequences of the SNPs identified by BEATRICE and SuSiE, respectively. Both approaches tag genes that involve APOE, TOMM40 [11], APOC1 [12], and PVRL2 [13], all of which have been previously associated with AD.

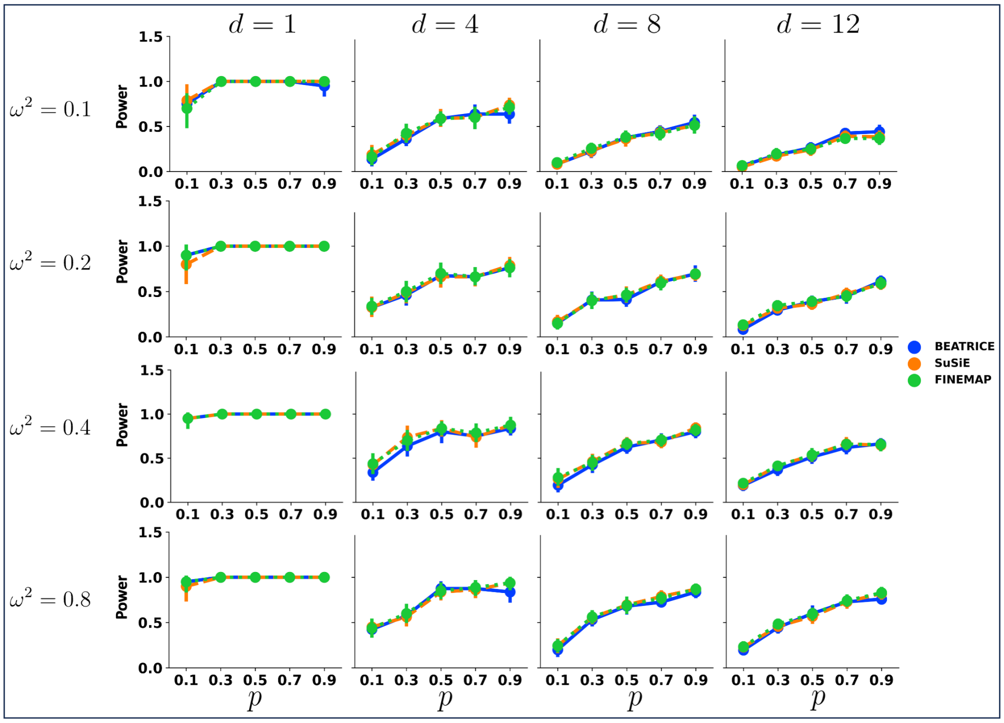


**Fig S10.** Power of the credible sets generated by three models across multiple causal variants $d = [1, 4, 8, 12]$, multiple SNP heritability $\omega^2 = [0.1, 0.2, 0.4, 0.8]$ and multiple infinitesimal effects from non-causal variants $p = [0.1, 0.3, 0.5, 0.7, 0.9]$. Each row and column corresponds to a specific value of $\omega^2$ and $d$, respectively. In each plot, the y-axis captures power, and the x-axis represents $p$.

However, only BEATRICE can pinpoint the APOE $\epsilon 2$ allele, which is commonly associated with the disease pathology of Alzheimer's disease. In conclusion, through this exploratory analysis, we show that BEATRICE can successfully parse complex LD regions and find putative causal factors in real-world datasets.

#### S1.18 Extending BEATRICE to Multiple Studies

BEATRICE has a simple and flexible design. Importantly, BEATRICE can easily incorporate priors based on the functional annotations of the variants. Formally, in the current setup, the prior over $\mathbf{c}$ is effectively constant, as captured by $p_0 = \frac{1}{m}$. We can integrate functional information [14] simply by modifying the distribution of $p_0$ across the variants. Thus, BEATRICE is a general-purpose tool for fine-mapping. Going one step further, a recent direction in fine-mapping is to aggregate data across multiple studies to identify causal variants [3,15]. Here, different LD matrices across studies help to refine the fine-mapping results. BEATRICE can be applied in this context as well by modifying Eq. (12) as

$$\mathcal{L}(\phi) = -\frac{1}{S\,L}\sum_{s=1}^{S}\sum_{l=1}^{L}\log\left(N\left(\mathbf{z}_s; 0, \mathbf{\Sigma}_{X_s} + \mathbf{\Sigma}_{X_s}\left(n\sigma^2\mathbf{\Sigma}_C^l(\phi)\right)\mathbf{\Sigma}_{X_s}\right)\right) + \sum_i \mathbf{p}_i \log\left(\frac{\mathbf{p}_i}{p_0}\right) + (1-\mathbf{p}_i)\log\left(\frac{1-\mathbf{p}_i}{1-p_0}\right) \quad (9)$$

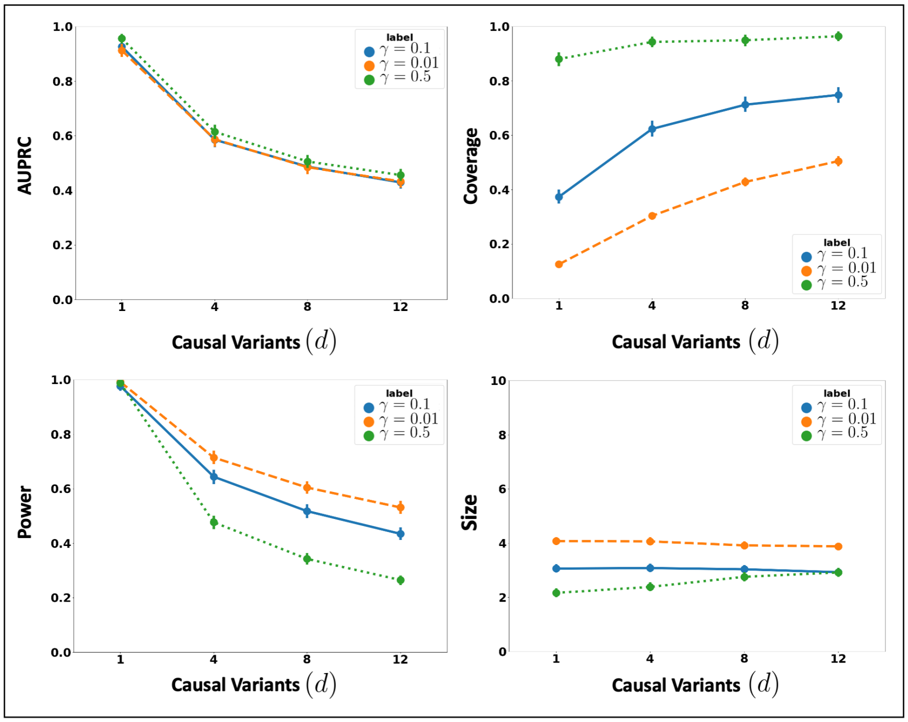


**Fig S11.** The performance metrics obtained by BEATRICE for three different values of $\gamma$ across varying numbers of causal variants. Along the x-axis, we plot the number of causal variants, and across the y-axis, we plot the mean and confidence interval (95%) of each metric. We calculate the mean by fixing $d$ to a specific value $d = d^*$ and sweep over all the noise settings where $d = d^*$.

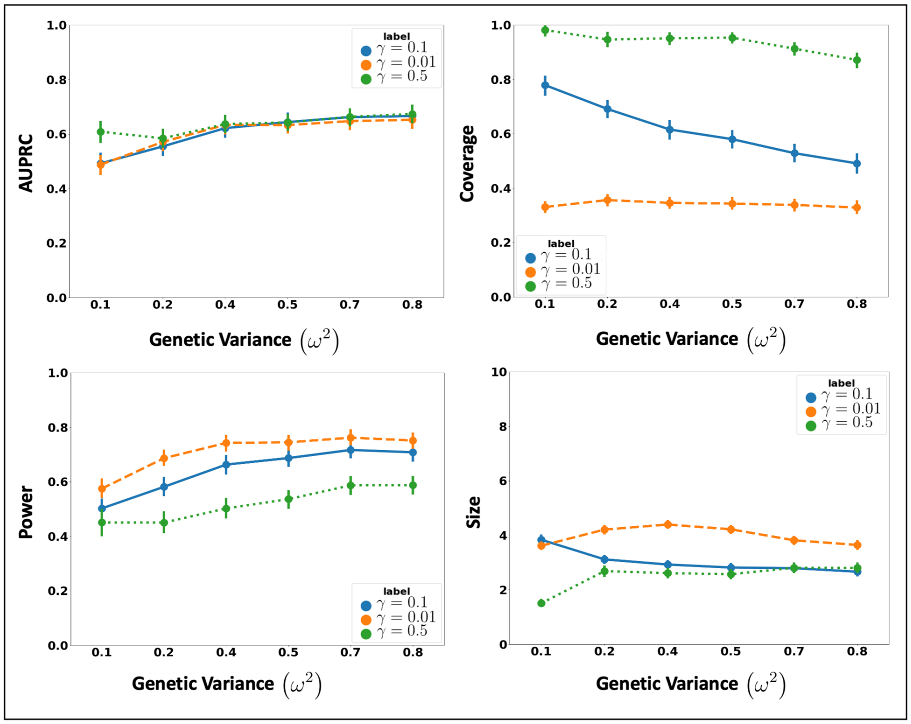


**Fig S12.** The performance metrics obtained by BEATRICE for three different values of $\gamma$ with increasing phenotype variance explained by genetics. Along the x-axis, we plot the variance explained by genetics ($\omega^2$), and across the y-axis, we plot the mean and confidence interval (95%) of each metric. We calculate the mean by fixing $\omega^2$ to a specific value $\omega = \omega^*$ and sweep over all the noise settings where $\omega = \omega^*$.

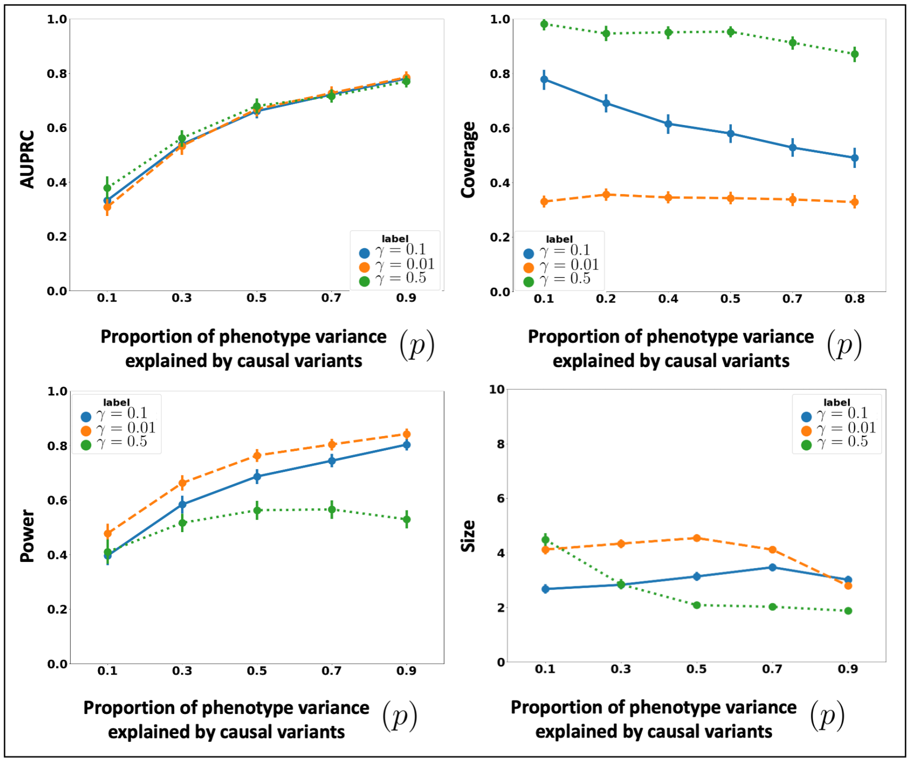


**Fig S13.** The performance metrics obtained by BEATRICE for three different values of $\gamma$ for multiple levels of noise introduced by non-causal variants. The noise level $(p)$ is explained by the variance ratio of non-causal variants vs. causal variants. Along the x-axis, we plot the noise level $(p)$; across the y-axis, we plot the mean and confidence interval (95%) of each metric. We calculate the mean by fixing $p$ to a specific value $p = p^*$ and sweep over all the noise settings where $p = p^*$.

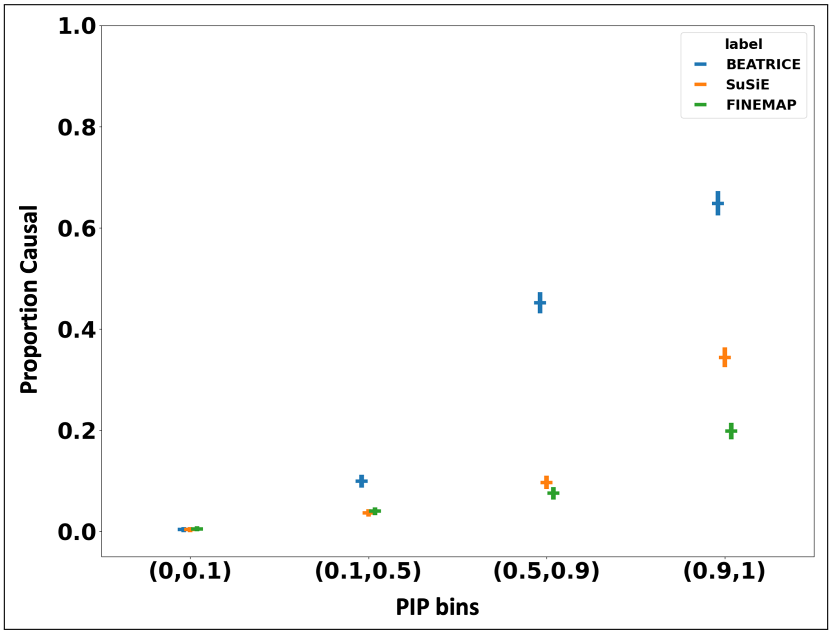


**Fig S14.** The calibration plot of the three fine-mapping methods aggregated across the 2400 simulation settings. The x-axis shows the PIP bins. The y-axis show the proportion of causal SNPs present within the PIP bin.

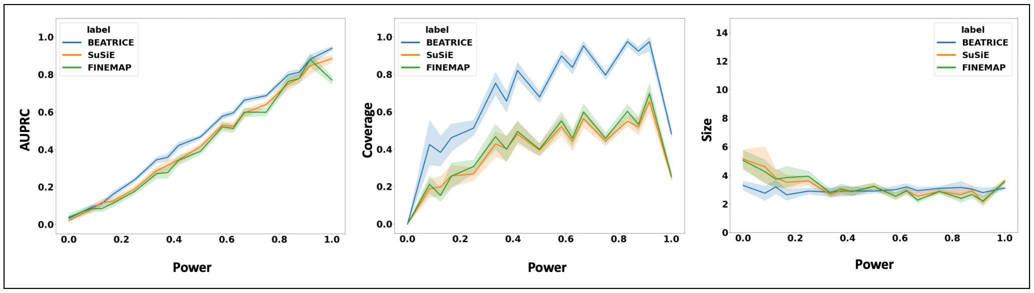


**Fig S15.** The change in AUPRC, coverage, size of credible set for constant power across 2400 simulation experiments. The shaded region represents the 95% confidence interval.

where $s$ denotes each separate study, $S$ is the total number of studies in the analysis, and $\mathbf{z}_s, \mathbf{\Sigma}_{X_s}$ are the summary statistics for each study. This same formulation can be used for multiple ancestries or traits. In this case, the "study" $s$ would correspond to an ancestry or trait. This proposed extension differs from the current formulation of BEATRICE on how we create the parameters $\mathbf{p}$ of the binary causal vectors. Currently, BEATRICE generates the paramters $\mathbf{p}$ from a single vector of $\mathbf{z}$ scores. The extension would now generate the parameters as a function of multiple $\mathbf{z}_s$ scores, corresponding to each study or ancestry, as $\mathbf{p} = f(\mathbf{z}_1, \dots, \mathbf{z}_S; \phi)$.

#### S1.19 Hyper-parameters of the Baseline Methods

We compare BEATRICE with SuSiE-v0.12.27, FINEMAP-v1.4.1, and CARMA-1.0.

**FINEMAP** This approach uses a stochastic shotgun search to identify causal configurations with non-negligible posterior probability. FINEMAP defines the neighborhood of a configuration at every step by deleting, changing or adding a causal

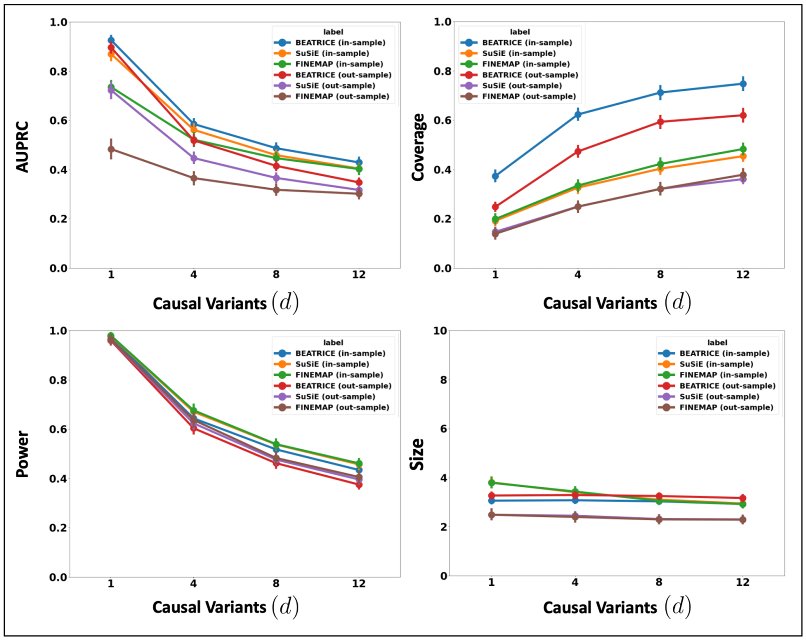


**Fig S16.** The performance metrics for the three methods across varying numbers of causal variants in the presence of both in-sample and out-of-sample LD matrices. Along the x-axis, we plot the number of causal variants, and across the y-axis, we plot the mean and confidence interval (95%) of each metric. We calculate the mean by fixing $d$ to a specific value $d = d^*$ and sweep over all the noise settings where $d = d^*$. In brackets, we mention whether that plot is generated from results obtained from in-sample or out-of-sample LD matrices.

variant from the current configuration. The next iteration samples from this neighborhood, thus reducing the exponential search space to a smaller high-probability region. Finally, the identified causal configurations are used to determine the posterior inclusion probabilities for each variant. The computationally efficient shotgun approach makes FINEMAP a viable tool for finemapping from multiple GWAS summary data in [16, 17]. We prune the credible sets of FINEMAP [18] via the approach used in [5] for this task.

We implement FINEMAP using the stochastic shotgun approach. During implementation, we fix the number of causal variants to 20 and the rest of the hyperparameters are fixed to default values, as described in http://christianbenner.com/

**SuSiE** [5, 19] introduced an iterative Bayesian selection approach for fine-mapping that represents the variant effect sizes as a sum of “single-effect” vectors. Each vector contains only one non-zero element, which represents the causal signal. In addition to finding causal variants, SuSiE provides a way to quantify the uncertainty of the causal variants locations via credible sets. SuSiE has also been used widely to find putative causal variants GWAS summary statistics [20, 21].

During the implementation of SuSiE, we provide the un-normalized effect sizes ($\beta$), the Standard Error (SE) of the effect sizes, the LD matrix, the phenotype variance, and the number of samples. Additionally, we fix the number of causal variants to 20 and we estimate the residual variance. The rest of the hyperparameters are fixed to default values, as described in
https://stephenslab.github.io/susieR/articles/finemapping_summary_statistics.html

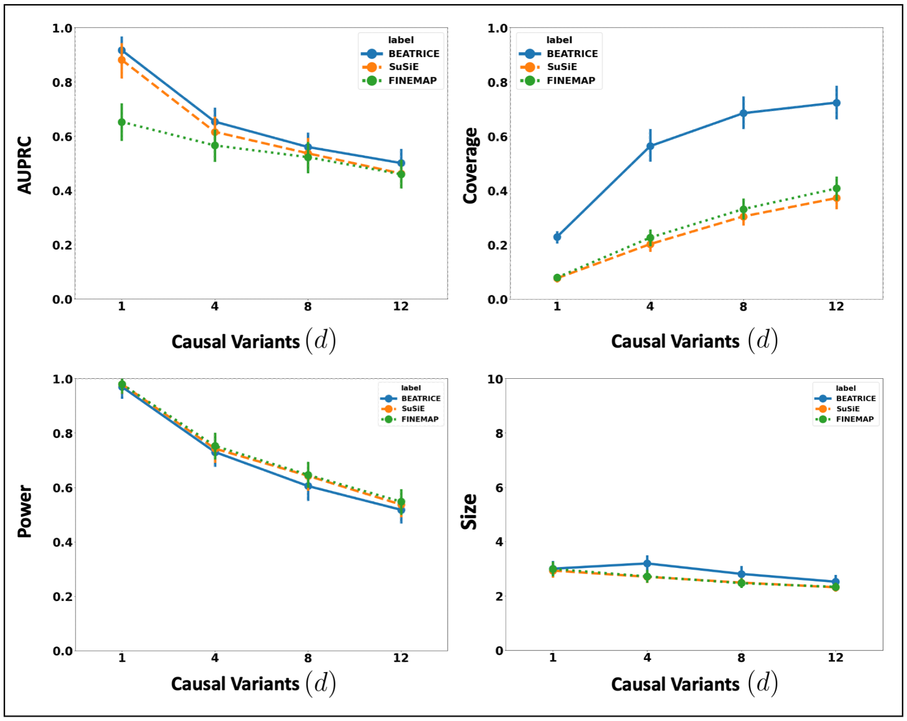


**Fig S17.** The performance metrics for the three methods across varying numbers of causal variants, while fixing the phenotypic variance explained by genetics at 0.5 and the variance explained by covariates at 0.2. Along the x-axis, we plot the number of causal variants, and across the y-axis, we plot the mean and confidence interval (95%) of each metric.

**CARMA** The recent work of [22] introduced a Bayesian approach for fine-mapping that assumes a spike-and-slab prior over the effect sizes. CARMA uses an MCMC sampling procedure to estimate the posterior distribution over the causal SNPs. In addition, the authors introduce a Bayesian hypothesis testing framework to prune out outliers in the presence of out-of-sample LD matrices and varying sample sizes across studies when computing the LD matrix.

We run CARMA in its default mode with spike-and-slab prior over the effect sizes as described in *https://github.com/ZikunY/CARMA/blob/master/CARMA_demo.pdf*

#### S1.20 Code Availability

We have compiled the code for BEATRICE and its dependencies into a docker image, which can be found at *https://github.com/sayangsep/Beatrice-Finemapping*. We have also provided installation instructions and a detailed description of the usage. The compact packaging will allow any user to directly download and run BEATRICE on their data. Namely, all the user must specify are a directory path to the summary statistics (i.e., z-scores), the LD matrix, and the number of subjects. Figure S25 shows the outputs generated by BEATRICE. The results are output in (1) a PDF document that displays the PIPs and corresponding credible sets, (2) a table with PIPs, (3) a text file with credible sets, and (4) a text file with the conditional inclusion probability of the variants within the credible sets. The user can also generate the neural network losses describe in Eq. (12) by adding a flag to the run command.

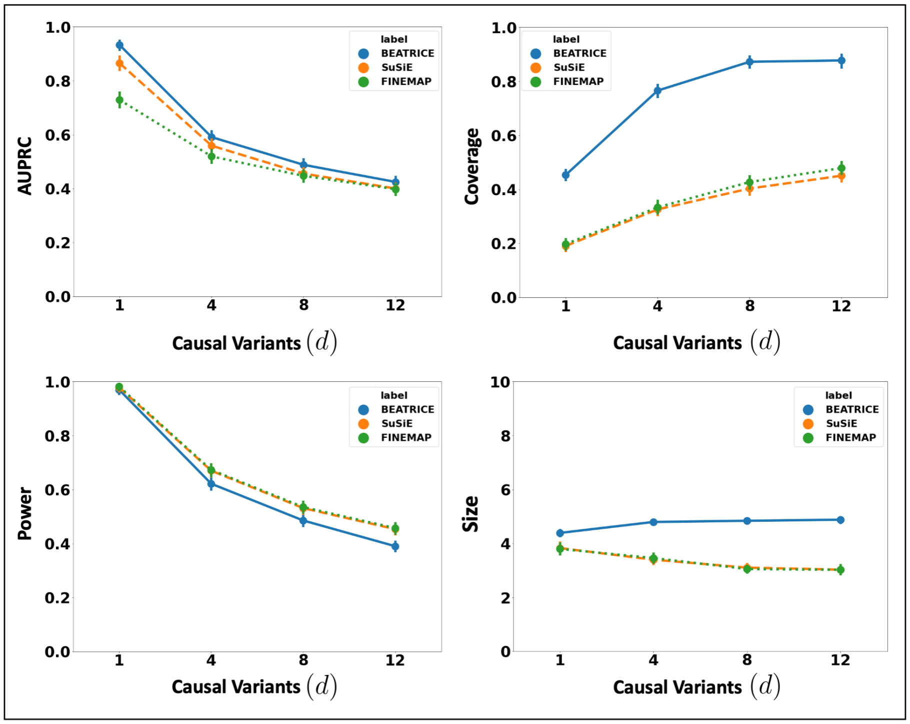


**Fig S18.** The performance metrics for the three methods across varying numbers of causal variants. Along the x-axis, we plot the number of causal variants, and across the y-axis, we plot the mean and confidence interval (95%) of each metric. We calculate the mean by fixing $d$ to a specific value $d = d^*$ and sweep over all the noise settings where $d = d^*$.

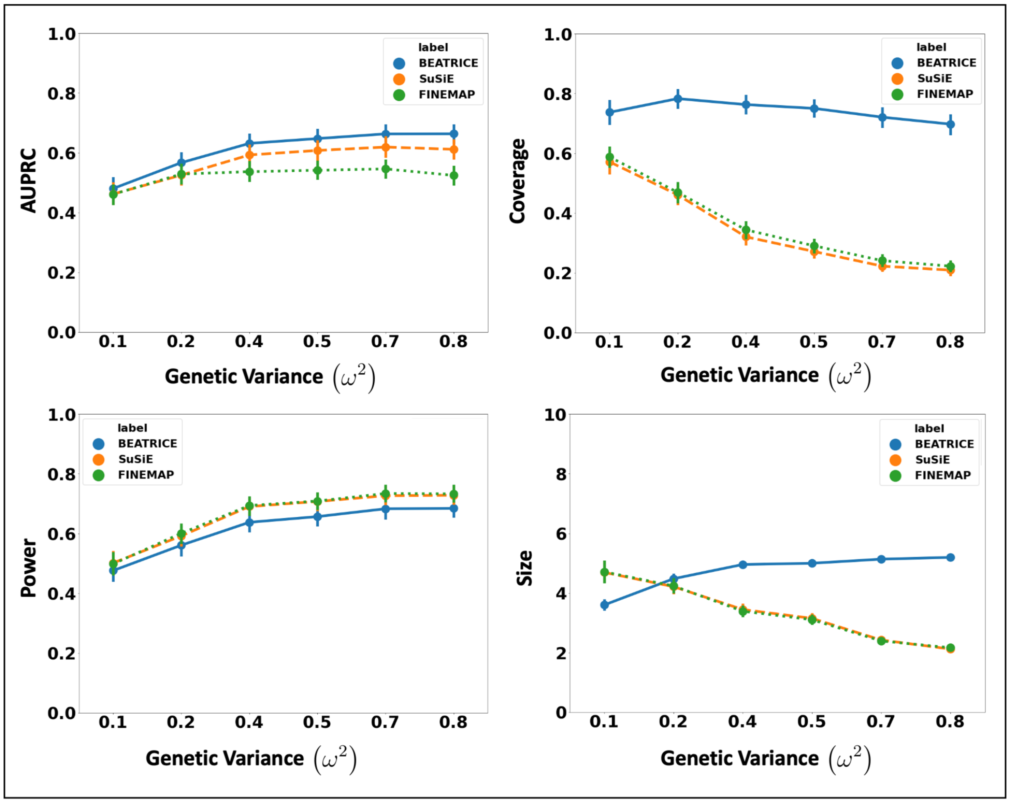


**Fig S19.** The performance metric for increasing phenotype variance explained by genetics. Along the x-axis, we plot the variance explained by genetics ($\omega^2$), and across the y-axis, we plot each metric's mean and confidence interval (95%). We calculate the mean by fixing $\omega^2$ to a specific value $\omega = \omega^*$ and sweep over all the noise settings where $\omega = \omega^*$.

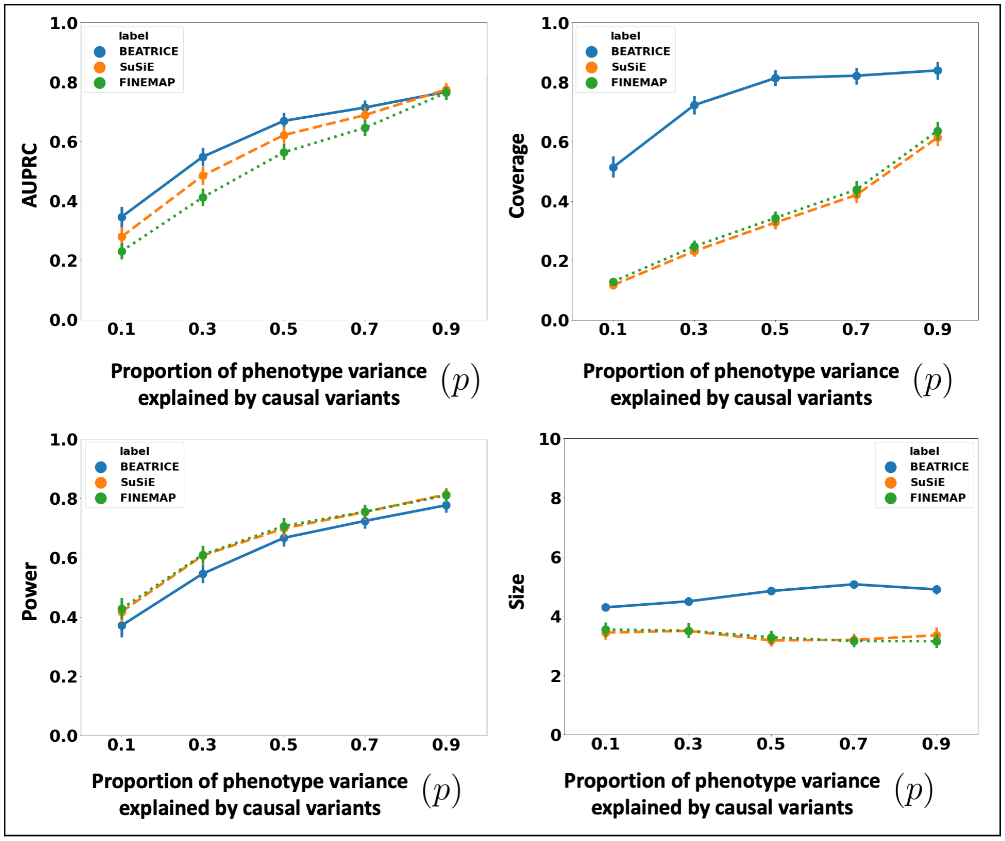


**Fig S20.** The performance metric for multiple levels of noise introduced by non-causal variants. The noise level ($p$) is explained by the variance ratio of non-causal variants vs. causal variants. Along the x-axis, we plot the noise level ($p$); across the y-axis, we plot each metric's mean and confidence interval (95%). We calculate the mean by fixing $p$ to a specific value $p = p^*$ and sweep over all the noise settings where $p = p^*$.

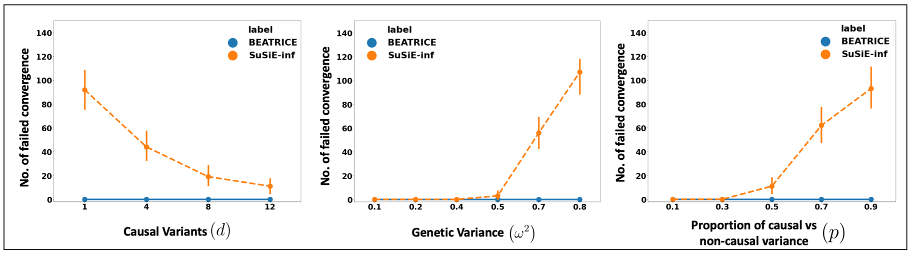


**Fig S21.** Number of non-convergent runs of SuSiE-inf, as compared to BEATRICE.

20. Albiñana C, et al. Genetic correlates of vitamin D-binding protein and 25-hydroxyvitamin D in neonatal dried blood spots. Nature Communications 2023 14:1. 2023;14(1):1–16. doi:10.1038/s41467-023-36392-5.

21. Li Y, , et al. Cross-ancestry genome-wide association study and systems-level integrative analyses implicate new risk genes and therapeutic targets for depression. medRxiv. 2023; p. 2023.02.24.23286411. doi:10.1101/2023.02.24.23286411.

22. Yang Z, et al. CARMA is a new Bayesian model for fine-mapping in genome-wide association meta-analyses. Nature Genetics 2023 55:6. 2023;55(6):1057–1065.

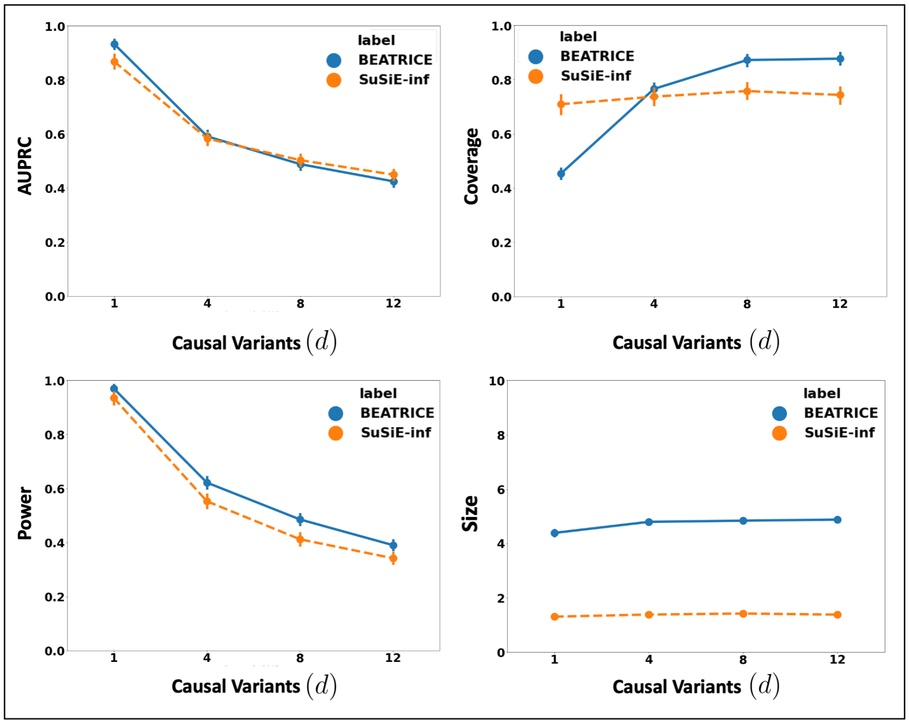


**Fig S22.** Performance metrics of BEATRICE and SuSiE-inf across varying numbers of causal variants. The performance of SuSiE-inf is calculated over the subset of simulation settings in which the algorithm converges; non-convergent settings are omitted from the analysis. The x-axis corresponds to the number of causal variants, and the y-axis plots the mean and confidence interval (95%) of each metric. We calculate the mean by fixing $d$ to a value $d = d^*$ and sweeping over all the noise settings where $d = d^*$.

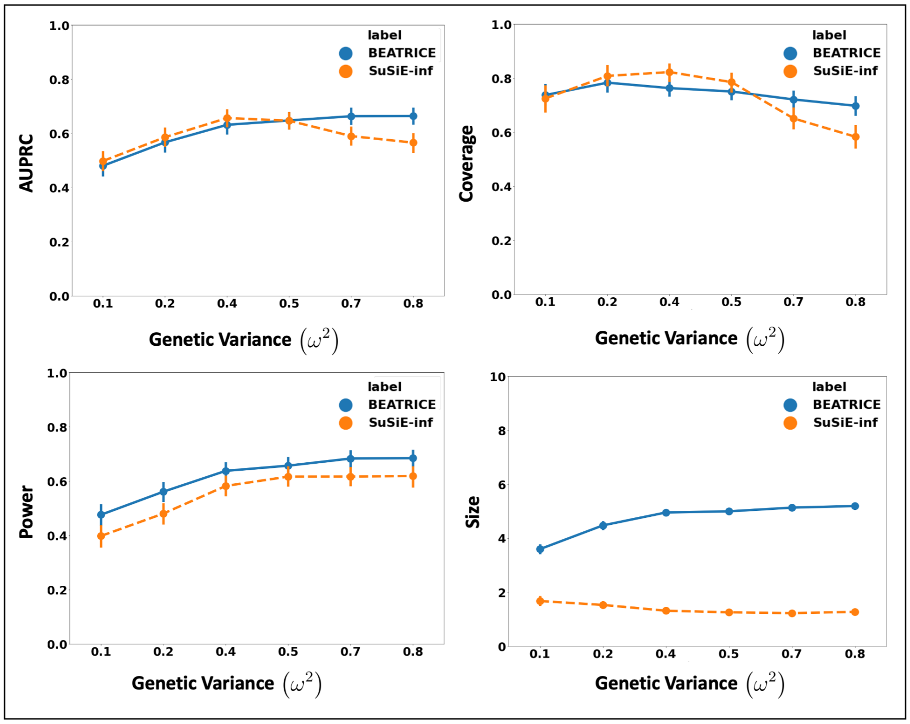


**Fig S23.** Performance metrics of BEATRICE and SuSiE-inf for increasing phenotype variance explained by genetics. The performance of SuSiE-inf is calculated over the subset of simulation settings in which the algorithm converges; non-convergent settings are omitted from the analysis. The x-axis corresponds to the variance explained by genetics ($\omega^2$), and the y-axis plots the mean and confidence interval (95%) of each metric. We calculate the mean by fixing $\omega^2$ to a value $\omega = \omega^*$ and sweeping over all the noise settings where $\omega = \omega^*$.

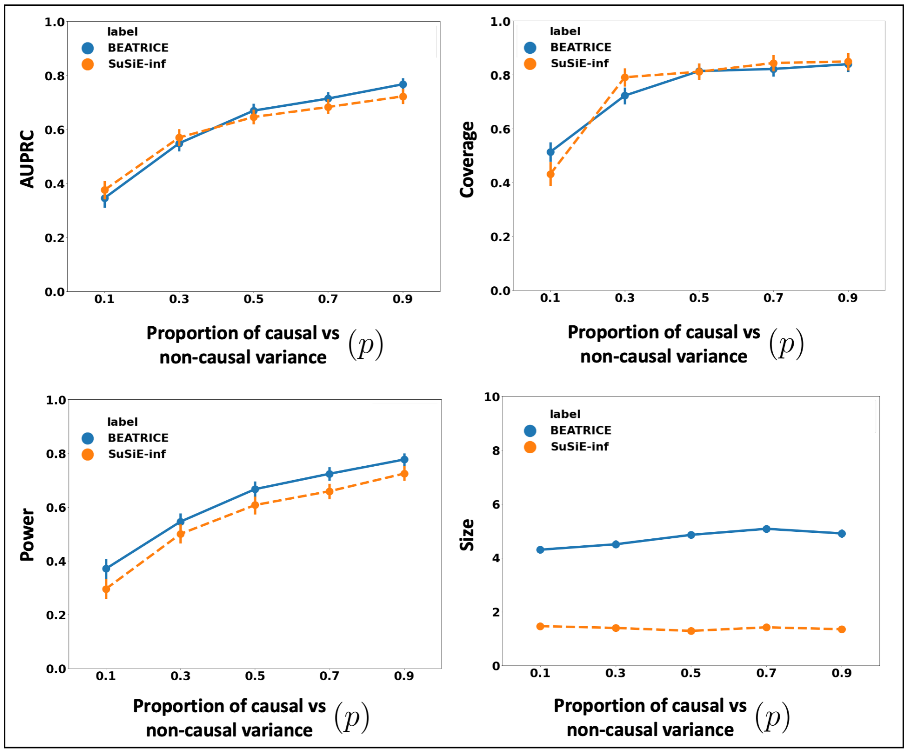


**Fig S24.** Performance metrics of BEATRICE and SuSiE-inf for multiple levels of noise introduced by non-causal variants. The performance of SuSiE-inf is calculated over the subset of simulation settings in which the algorithm converges; non-convergent settings are omitted from the analysis. The x-axis corresponds to the noise level ($p$), and the y-axis plots the mean and confidence interval (95%) of each metric. We calculate the mean by fixing $p$ to a value $p = p^*$ and sweeping over all the noise settings where $p = p^*$.

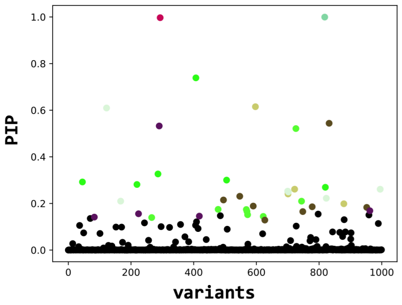


(a)

| Variant index | pip | Variant names |
|---|---|---|
| 818 | 0.99 | rs818 |
| 293 | 0.99 | rs293 |
| 407 | 0.73 | rs407 |
| 597 | 0.61 | rs597 |
| 122 | 0.60 | rs122 |

(b)

```
818
293
407 505 45 286 219 820
122 995 700 823 702 167
597 701 879 722
```

(c)

```
1.0
0.996
0.592 0.088 0.087 0.084 0.081 0.079
0.583 0.09 0.088 0.088 0.086 0.081
0.699 0.087 0.084 0.084
```

(d)

**Fig S25.** Overview of the outputs generated by BEATRICE. (a) The PIPs are displayed and color coded by their assignment to credible sets. (b) A table with the PIPs and the corresponding name of the variants. (c) A text file with the credible sets. Here each row represent a credible set and the entries are indices of the variants present in the credible set. The first column of each row represents the key index. (d) The conditional inclusion probability of each of the credible variants given all the key variants. The calculations can be found in **Section S1.3** of the Supplements.
